# p300/CBP-catalyzed acetylation safeguards minor zygotic genome activation via activating DUX

**DOI:** 10.1101/2024.11.27.625783

**Authors:** Lieying Xiao, Hao Jin, Yanna Dang, Panpan Zhao, Shuang Li, Yan Shi, Shaohua Wang, Kun Zhang

## Abstract

Zygotic genome activation (ZGA), the first transcriptional event at the onset of life, is crucial for early embryonic development. However, the mechanisms that initiate and regulate ZGA in mammals remain poorly understood. In this study, we demonstrate that p300/CBP-catalyzed acetylation plays a vital role in regulating ZGA. Inhibition of p300/CBP acetyltransferase activity results in impaired minor ZGA and a subsequent 2-cell arrest. By profiling the dynamic landscape of p300 during the ZGA stage, we elucidate the distinct patterns of p300 establishment in promoter and enhancer regions. Moreover, p300 recruits RNA polymerase (Pol II) to pre-configure in the promoter regions of ZGA genes, and orchestrates the gradual establishment of enhancer activity during the early 2-cell stage to activate ZGA. Furthermore, p300 enhances minor ZGA by promoting the expression of DUX. Restoring DUX levels, which are diminished due to p300/CBP inhibition, rescues ZGA and embryonic development by re-establishing Pol II enrichment on minor ZGA genes. Overall, our findings elucidate the mechanism by which p300/CBP regulates ZGA through acetylation and activation of the key transcription factor DUX.

**Highlights:**

- p300/CBP plays a stage-specific role during preimplantation development phase.
- p300 displays distinctive dynamic changes in promoter and enhancer regions during ZGA stage.
- p300/CBP-catalyzed acetylation facilitates the pre-configuration and elongation of Pol II at ZGA gene regions.
- DUX recruits p300 and Pol II to activate minor ZGA genes independently of p300’s catalytic activity.

## Introduction

After fertilization, terminally differentiated gametes fuse to form a totipotent zygote, capable of generating an entire organism^1^. The zygote undergoes pre-implantation development to form blastocysts, which involves a series of significant biological events, including zygotic genome activation (ZGA), followed by the first cell-fate decision and lineage specification. ZGA, the first transcriptional event, is accompanied by the degradation of maternal RNA and proteins and essential for embryonic development^2-4^. In mice, ZGA initiates during the S phase of the zygote and the G1 phase of the early 2-cell (E2C) stage, resulting in the activation of only a limited number of genes and exhibiting a low level of promiscuous transcription, a process termed minor ZGA. A substantial increase in transcription occurs at the late 2-cell (L2C) stage, designated as major ZGA. Timely regulation of both minor and major ZGA is crucial for early embryo development and requires a functional and precise regulatory network^3-5^. Despite its importance, the molecular regulation of ZGA remains poorly understood.

Key transcription factors (TFs) are often considered the primary drivers in the activation of specific gene transcription and the subsequent triggering of ZGA^6,7^. In mice, the OBOX family provides a prominent example. Maternal or zygotic TF OBOXs play a pivotal role in the recruitment of Pol II to the promoter regions of ZGA genes^5^. Other TFs, including DUX^8-10^ and NFYA^11^, have also been shown to regulate ZGA genes. However, TFs alone are not sufficient for the activation of specific gene expression. TFs that are bound to specific sequences at enhancers require the involvement of mediators or other coactivators to bridge them to promoters, thereby enabling the recruitment of the transcription machinery^13,14^. The histone acetyltransferase (HAT) paralogs p300 and CBP (collectively referred to as p300/CBP) are essential transcriptional co-activators that act as scaffolds to mediate the binding of TFs and Pol II^15,16^. p300/CBP are primarily involved in enhancer-dependent transcription regulation through their acetylation activity. H3K27ac, catalyzed by p300/CBP, is widely recognized as a hallmark of active promoters and enhancers^17,18^. Furthermore, super-enhancers (SEs) are characterized by significant levels of p300 and H3K27ac^19,20^.

p300/CBP plays a pivotal role in vertebrate development. In mice, homozygous knockout of *Ep300*^-/-^ or *Crebbp*^-/-^ results in post-implantation lethality, and even double heterozygous knockout mice (*Ep300*^+/-^ / *Crebbp*^+/-^) die during embryonic development^21,22^. This suggests that p300 and CBP have overlapping functions in embryonic development, and that mouse embryos are particularly susceptible to fluctuations in the dosage of p300/CBP, exhibiting a dose-dependent effect. Recent studies have demonstrated that p300/CBP plays a role in the promotion of ZGA. In *Drosophila*, the CBP homologue Nejire has been demonstrated to regulate the expression of ZGA genes, and its catalytic activity has been shown to be dispensable for the proper functioning of ZGA^23^. In *zebrafish*, maternal knockdown of *ep300b* and *crebbpa*/*crebbpb* has been observed to result in a decline in H3K27ac and the downregulation of ZGA-related genes, ultimately leading to developmental arrest at the ZGA stage^24^. Overexpression of p300 and Brd4 in zygotes causes an early accumulation of H3K27ac at ZGA-related genes, promoting premature ZGA initiation^25^. In mice, it has recently been determined that the HAT activity of p300/CBP is crucial for mouse ZGA and early embryonic development. Inhibition of p300/CBP results in the failure of ZGA and developmental arrest^26^. However, the precise mechanisms by which p300/CBP participate in the transcription initiation and elongation of ZGA genes remain unclear.

Here, we conducted a comprehensive investigation into the functions and mechanisms of p300/CBP-catalyzed acetylation in ZGA. Our findings reveal that p300/CBP-catalyzed acetylation is a crucial driver for the activation of minor ZGA genes, as well as for early mouse embryonic development. By profiling the dynamic landscape of p300 during the mouse ZGA stage, we observed that p300 recruits Pol II to pre-configure at the promoters of major ZGA genes during the early 2-cell (E2C) stage. Additionally, we noted a stepwise establishment of enhancer activity and the enhancer regulatory network from the E2C to the late 2-cell (L2C) stage. Furthermore, DUX emerges as a key TF regulating the initiation of minor ZGA transcription and Pol II pre-configuration.

## Results

### p300/CBP-catalyzed acetylation is critical for minor ZGA and embryogenesis beyond the 2-cell stage in mice

Initially, we examined the expression levels of p300/CBP in mouse oocytes and preimplantation embryos by analyzing published data^27^ and conducting immunostaining. Results revealed maternal expression of *Ep300* and *Crebbp*, with significant activation occurring at the 2-cell stage (Figure S1A). Furthermore, high protein levels of p300/CBP were sustained throughout germinal vesicle (GV) oocytes and the entire pre-implantation developmental stage (Figure S1B), suggesting that p300/CBP may play a role during maternal-to-zygotic transition (MZT).

To explore the role of p300/CBP during pre-implantation development, we utilized a highly selective inhibitor, A485^28^, to inhibit the catalytic activities of p300/CBP in a series of crucial cellular events, including ZGA and cell fate commitment (post-ZGA) (Figure 1A; see Methods). The inhibition by A485 was reversible and selective, as evidenced by the elimination and subsequent restoration of its target histone acetylations^18^, such as H3K27ac, H3K18ac, and H2BK5ac, upon A485 treatment and subsequent washing (Figure S1C). Notably, other histone acetylations, such as H3K14ac and H4K16ac, remained unaffected by A485 treatment (Figure S1D).

**Figure 1.**
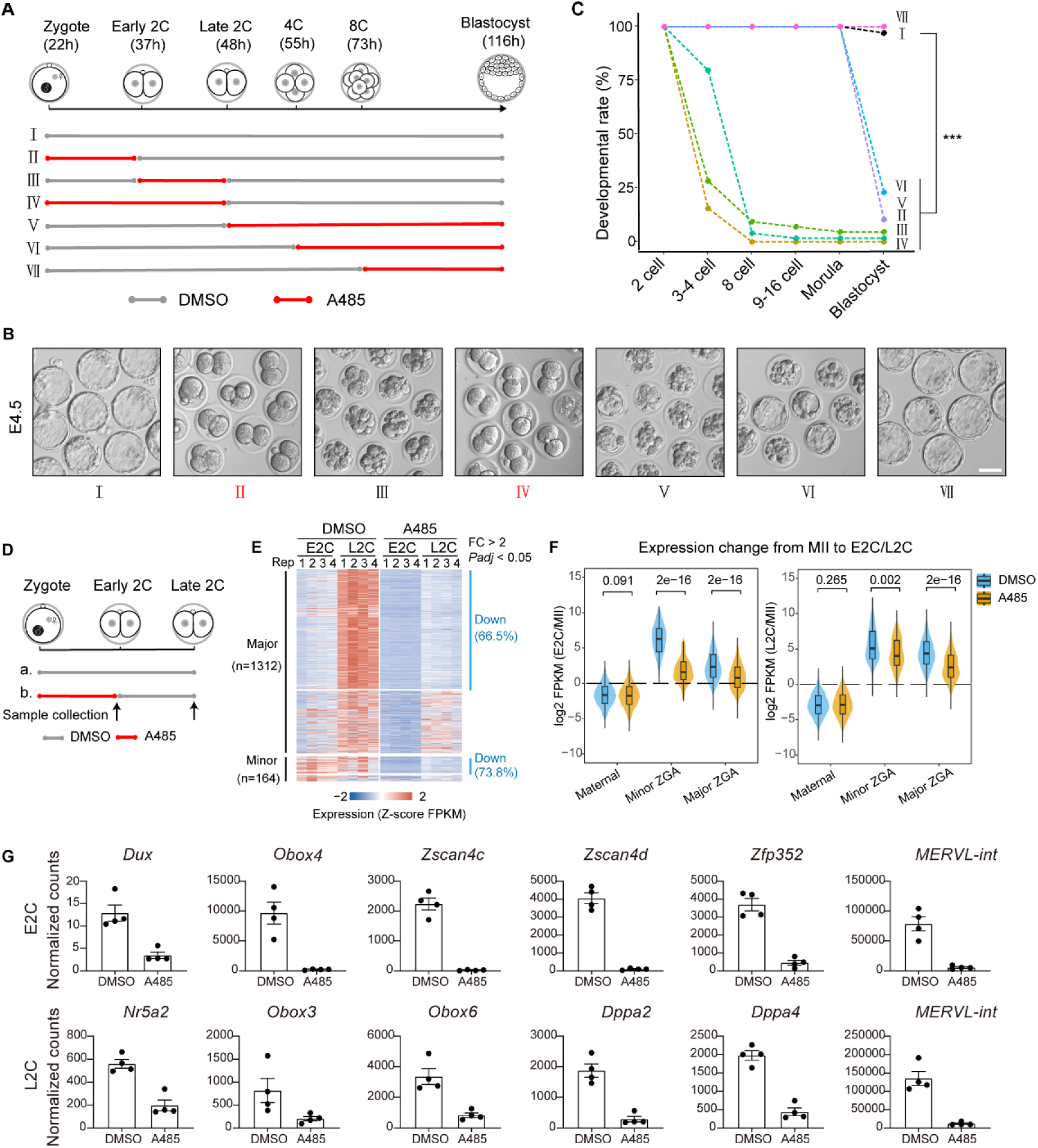
p300/CBP inhibition during minor ZGA leads to ZGA failure and 2-cell arrest. (A) Schematic showing p300/CBP inhibition by A485 treatment during minor ZGA (Ⅱ), major ZGA (Ⅲ), whole ZGA (Ⅳ), and from late 2-cell (Ⅴ), 4-cell (Ⅵ) and 8-cell (Ⅶ) and control group (I). (B) The phenotype showing the embryo morphology at E4.5 in different groups. Scale bar, 50 μm. (C) The developmental rate of embryos treated with A485 versus control (DMSO). *** *P* < 0.001. (D) Schematic showing the sample collection for RNA-seq with p300/CBP inhibition during minor ZGA at early 2-cell (E2C) and late 2-cell (L2C) stage. (E) Heatmap showing the minor and major ZGA gene expression in control (DMSO) and A485-treated embryos (4 biological replicates for E2C and L2C). n, gene number. FC, fold change; *Padj*, adjusted *P*-value. (F) Violin plot showing the expression changes of maternal, minor ZGA and major ZGA genes from MII oocytes to E2C or L2C in control (DMSO) and A485-treated embryos. Center line, median; box, 25th and 75th percentiles; whiskers, 1.5 × IQR. (G) Bar charts showing minor and major ZGA gene expression in control (DMSO) and A485-treated embryos (4 biological replicates for E2C and L2C). Data are mean ± S.E.M. *** *Padj* < 0.001.

Interestingly, the transient inhibition of p300/CBP during major ZGA resulted in embryonic arrest before compaction (Figures 1B and 1C, III), while transient inhibition during minor ZGA or throughout ZGA led to developmental arrest at the 2-cell stage (Figures 1B and 1C, II, IV). These phenotypes highlight the critical role of p300/CBP in ZGA and preimplantation embryo development, particularly its indispensable function in minor ZGA for development beyond the 2-cell stage. Moreover, embryos with transient inhibition of p300/CBP from the L2C or 4-cell (4C) stage to the blastocyst stage exhibited compromised blastocyst development (Figures 1B and 1C, V, VI), indicating that the catalytic activities of p300/CBP are also essential for post-ZGA development. However, p300/CBP is not essential during the 8-cell to blastocyst stages, as A485 treatment did not affect blastocyst development (Figures 1B and 1C, VII). These results indicate that the acetyltransferase activity of p300/CBP is critical for ZGA and preimplantation embryo development.

Previous research has shown that the activation of minor ZGA is critical for major ZGA and early embryo development^4^. Similarly, our findings revealed that impaired catalytic activity of p300/CBP specifically during minor ZGA resulted in 2-cell arrest, mirroring the outcomes observed when p300/CBP activity was blocked throughout the entire ZGA (Figures 1B and 1C). These results suggest that the role of p300/CBP in activating minor ZGA is crucial for both major ZGA and preimplantation embryo development. To further explore the role of p300/CBP during the minor ZGA stage, we aimed to determine how inhibition of p300/CBP during this phase led to 2-cell embryo arrest. We performed RNA sequencing on E2C and L2C embryos treated with DMSO and A485 to assess whether ZGA was affected (Figures 1D and S2A; Table S1). As expected, transcriptome analysis revealed a notable number of differentially expressed genes (DEGs) and differentially expressed repetitive elements in E2C (1719 and 47) and L2C (8354 and 270) (Figures S2B and S2C). MERVL repetitive elements, which are transiently upregulated during ZGA and are required for preimplantation embryo development^29^, were downregulated in both E2C and L2C (Figure 1G). The downregulated genes in these stages were predominantly involved in ribosome biogenesis, mRNA splicing, and the cell cycle (Figure S2D), indicating compromised ZGA.

In the E2C stage, 73.8% (121 out of 164) of minor ZGA genes were downregulated in the A485 group (Figures 1E and S2E; Table S2), including the TFs *Dux, Obox4*, and *Zscan4s* (Figure 1G). At the L2C stage, A485 treatment led to a significant decrease in major ZGA genes (860 out of 1312, 65.5%; Figures 1E and S2E; Table S3), including TFs *Obox3/6, Dppa2/4,* and *Nr5a2* (Figure 1G). The defects in ZGA were not due to developmental delays, as evidenced by the prompt global clearance of maternal transcripts (Figures 1F and S2F). Moreover, p300/CBP activity is essential for its own expression, as indicated by the decreased *Ep300* mRNA levels following A485 treatment at the L2C stage (Figure S2C). However, the abundant maternal p300 and CBP proteins can compensate for the reduction in zygotic p300 and CBP, mitigating the impact of this decrease (Figure S2G).

Additionally, a subset of genes exhibited ectopic transcription (Figure S2H), characterized by promiscuous transcription at the L2C stage with abundant intronic reads, and there was also a decrease in the expression of the histone demethylase *Kdm5b* along with impaired demethylation of H3K4me3 (Figures S2I and S2J), resembling the effects observed at the L2C stage after inhibiting minor ZGA transcription with DRB^3,4^. These findings suggest a failure in the transition from minor to major ZGA following A485 treatment. In sum, our results demonstrate that the acetyltransferase activity of p300/CBP during minor ZGA is critical for the timely onset of both minor and major ZGA.

### p300 is pre-configured at major ZGA genes prior to the late 2 cell stage

Given the high degree of homology between p300 and CBP^30^, we focused exclusively on p300 in the subsequent analyses. To elucidate the mechanisms by which p300/CBP regulates ZGA, we employed Stacc-seq^3^, a method analogous to CUT&Tag, which has been shown to be effective in analyzing limited cell counts. This was used to examine p300 distribution in control (considered wild-type, WT) and A485-treated L1C, E2C, and L2C embryos (Figure 2A). The Stacc-seq data for p300 demonstrated high reproducibility across replicates and revealed distinct binding patterns among different embryonic stages and treatments (Figures S3A and S3B).

**Figure 2.**
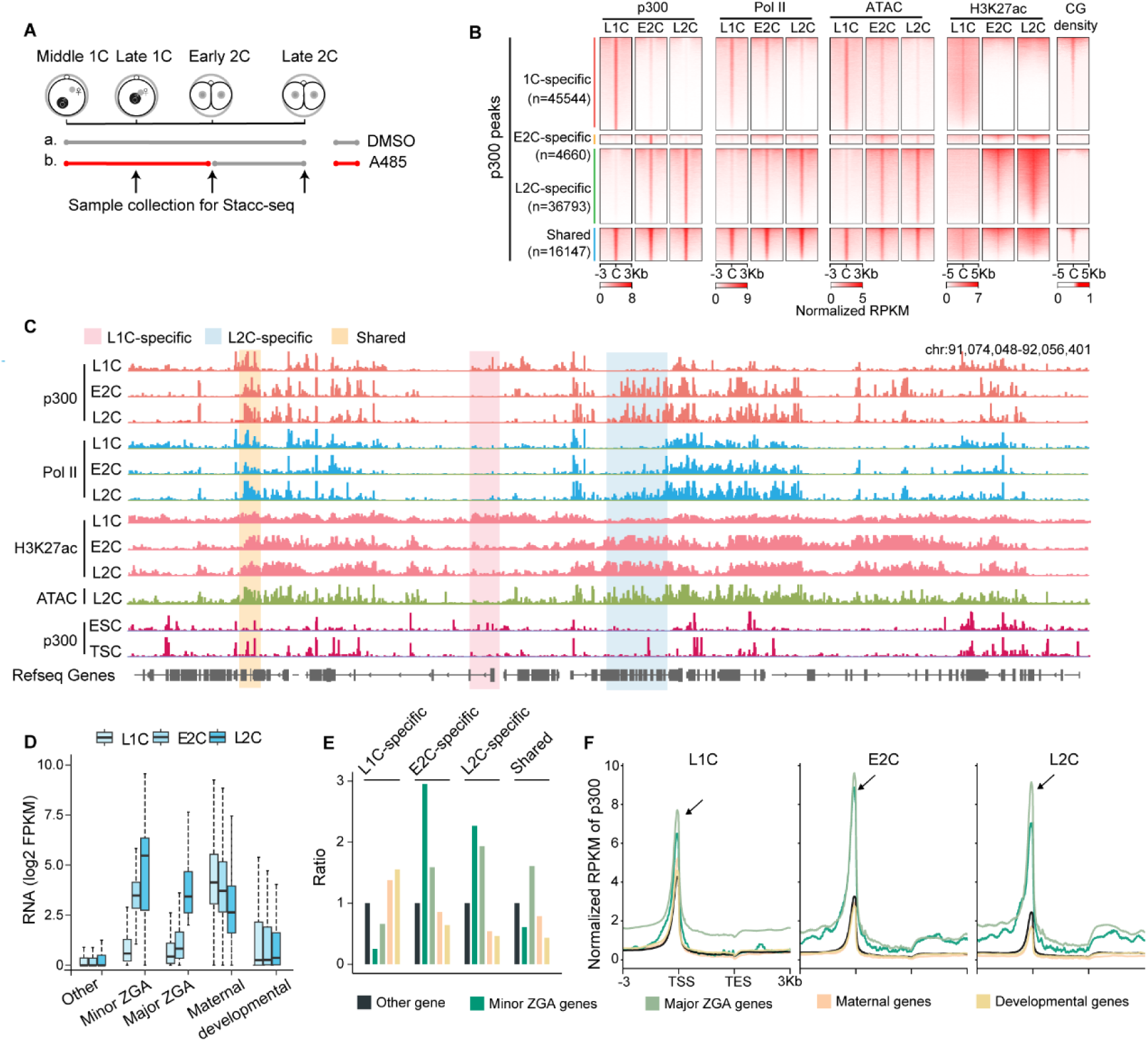
p300 is pre-configured at major ZGA genes prior to L2C. (A) Schematic showing the sample collection for Stacc-seq with p300/CBP inhibition during minor ZGA at late 1-cell (L1C), early 2-cell (E2C) and late 2-cell (L2C) stages. (B) Heatmaps showing p300 binding (2 biological replicates), Pol II binding, chromatin accessibility (ATAC-seq), H3K27ac (Z-score normalized) and CpG density at 1C-speific, E2C-specific, L2C-specific and shared p300 peaks in mouse embryos. Pol II Stacc-seq dataset (GSE135457), ATAC-seq dataset for L1C (GSE207222), H3K27ac CUT&RUN dataset (GSE207222) were used. (C) IGV browser snapshots showing p300, Pol II, H3K27ac enrichment and chromatin accessibility (ATAC-seq) in 1C, E2C, L2C embryos, p300 enrichment in ESCs (embryonic stem cells) and TSCs (trophoblast stem cell). Pol II Stacc-seq dataset (GSE135457), H3K27ac CUT&RUN dataset (GSE207222), p300 ChIP-seq data for ESCs and TSCs ((GSE110950)) were used. (D) Boxplot showing the expression (FPKM) of minor ZGA, major ZGA, maternal, developmental and other genes at individual stages. Center line, median; box, 25th and 75th percentiles; whiskers, 1.5 × IQR. (E) Histograms showing the proportion of minor ZGA, major ZGA, maternal and developmental genes occupied by promoter (around TSS < 1 kb) p300 peaks in 1C-speific, E2C-specific, L2C-specific and shared p300-binding regions. The expressed genes serve as a control. (F) Line chart showing the enrichment of p300 (Z-score normalized RPKM) in the five classes of genes at individual stages. TSS, transcription start site. TES, transcription end site.

Initially, we examined the distribution of p300 during the ZGA stage in WT embryos. As anticipated, the majority of p300-binding sites were located distally from the transcriptional start sites (TSSs) of well-annotated genes (Figure S3C). Notably, compared to other features, promoters exhibited higher occupancy scores for p300, which contrasts with observations in stem cells, where enrichment is uniformly distributed across all features (Figure S3D). This suggests that promoter-associated p300 play a critical role during the ZGA stage. Additionally, p300 preferentially occupied certain 2-cell-specific repeats, such as Alu, B2, and ERVL (Figure S3E).

The global distribution of p300 showed distinct stage-specific characteristics (Table S4). Notably, L1C-specific p300 experienced a significant decrease, while L2C-specific p300 exhibited a dramatic increase from E2C (Figures 2B and 2C). Compared to the other two classes, L1C-specific and stage-shared peaks demonstrated a greater tendency to be enriched in the promoter regions (20% and 43%, respectively), with these promoters being CG-rich (Figures S3F and 2B). Additionally, promoter-bound p300 displayed specificity for different gene categories: maternal genes, minor ZGA genes, major ZGA genes, and developmental genes (Figure 2D; Table S5; see Methods). L1C-specific p300 was associated with maternal and developmental genes enriched in multicellular organism development and cell differentiation (Figures 2E and S3G). In contrast, E2C-specific p300 bound to minor ZGA genes involved in protein import into nucleus and DNA replication initiation, which are pertinent to the G1 phase in E2C (Figures 2E and S3G). Both L2C-specific and stage-shared p300 preferentially associated with ZGA genes enriched in fundamental transcription pathways, such as ribosome biogenesis and RNA splicing (Figures 2E and S3G). These results indicate that p300 is initially enriched on ZGA and developmental genes with high-CG promoters during the L1C stage. Subsequently, at the E2C stage, p300 dissociates from developmental gene regions and begins to enrich on ZGA genes characterized by low-CG promoters.

To further investigate the regulatory relationship between p300 and ZGA genes, we mapped the enrichment dynamics of p300 on the promoters of four gene classes (Figures 2D and 2F). Interestingly, at the L1C stage, p300 exhibited similar levels of enrichment across all four gene classes (Figure 2F). However, p300 began to show increased enrichment specifically at ZGA genes during the E2C stage, maintaining a high level until the L2C stage. Concurrently, there was a dramatic decrease in p300 binding at maternal and developmental genes during E2C, with this decrease persisting until L2C (Figure 2F). These findings demonstrate a pre-loading of p300 on ZGA genes at the E2C stage, akin to the pre-configuration previously observed in Pol II (Figure 2B)^3^. This suggests that p300 facilitate the pre-configuration of Pol II on ZGA genes. Consistently, H3K27ac and chromatin accessibility exhibited similar dynamic changes alongside p300 and Pol II (Figures 2B and 2C). In summary, our data suggest that p300 and H3K27ac are pre-configured to an intermediate state prior to major ZGA.

### p300/CBP-catalyzed acetylation enhances the pre-configuration of p300 and pol II, facilitating pol II elongation at ZGA genes

To investigate the relationship between p300 and Pol II pre-configuration in ZGA, we focused on the specific binding of p300 and Pol II in 2C embryos. Approximately 90% of p300 peaks co-localized with Pol II in E2C and L2C embryos, while around 70% and 56% of Pol II peaks were unique to Pol II (Figures 3A and S4A). Among the promoter-bound Pol II peaks, those shared with p300 exhibited higher Pol II occupancy, and the associated genes showed elevated transcription levels (Figures 3A, 3B, S4B). Moreover, compared to Pol II-specific peaks, the shared peaks were more likely to be enriched in major ZGA gene regions during the E2C and L2C stages (Figures 3C and S4C). These findings indicate that p300 facilitates Pol II binding at ZGA genes during the 2C stage, correlating with the active transcription of mouse ZGA genes.

**Figure 3.**
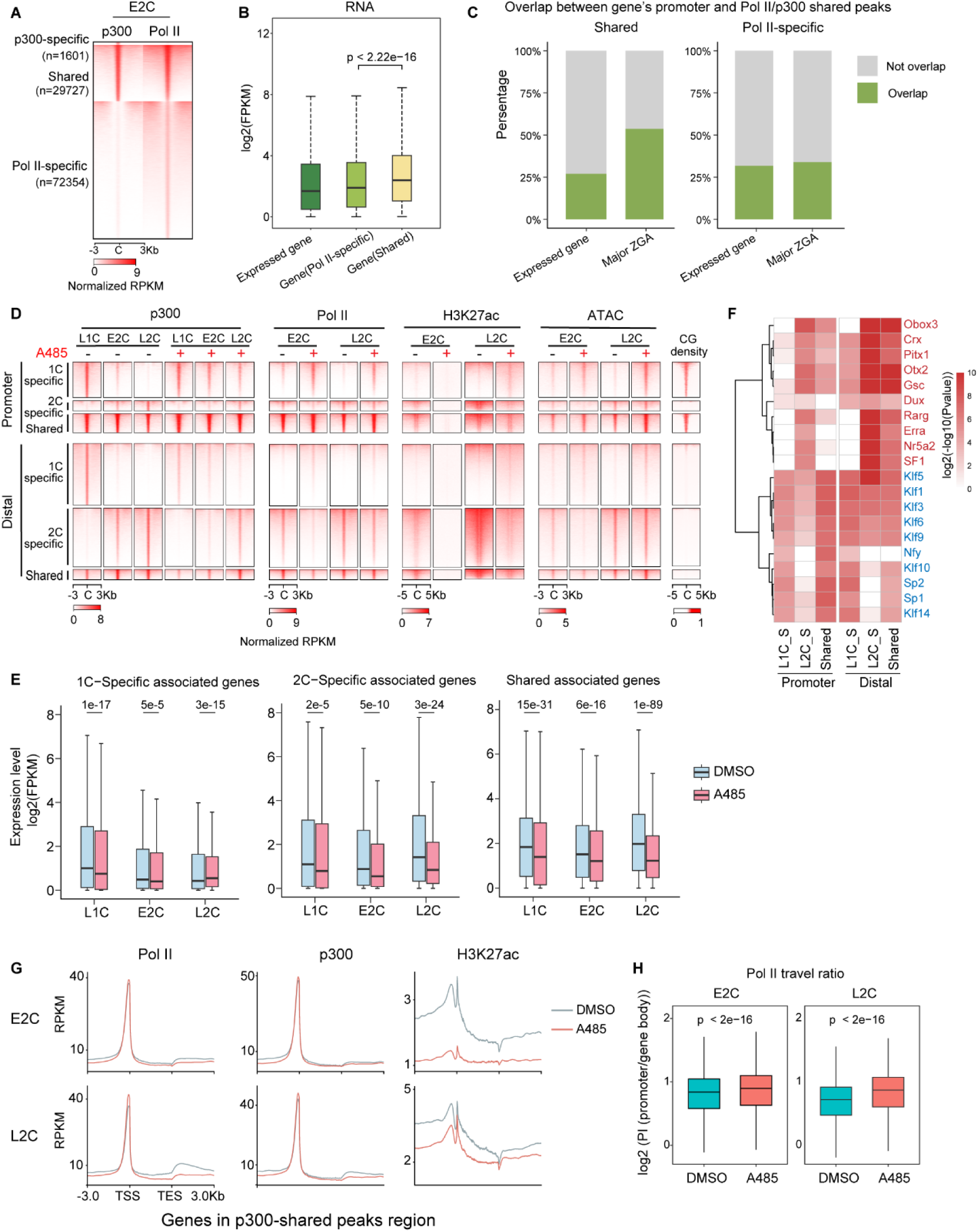
A485 results in the occurrence of pre-configuration defects in both p300 and Pol II, as well as the failure of Pol II elongation. (A) Heatmaps illustrating the specific and shared enrichment of promoter p300 and Pol II (GSE135457) at promoter regions in untreated E2C embryos. (B) Boxplot illustrating the expression levels of genes associated with different types of promoter peaks in E2C embryos. The p-values are indicated using a two-sided Wilcoxon rank-sum test. Center line, median; box, 25th and 75th percentiles; whiskers, 1.5 × IQR. (C) Histograms illustrating the proportion of expressed genes and major ZGA genes associated with shared peaks (left) and Pol II-specific peaks (right) in E2C embryos. The proportion of expressed genes and major ZGA genes is accounted for by the overlapping and non-overlapping. (D) Heatmaps illustrating p300 binding, Pol II binding, H3K27ac, chromatin accessibility (ATAC-seq) (Z-score normalized) and CpG density at 1C-specific, 2C-specific and shared p300 peaks in control (DMSO) (-) and A485-treated (+) embryos. (E) Boxplots illustrating the expression levels of genes associated with different regions from the L1C to L2C stages in the DMSO and A485 groups. Center line, median; box, 25th and 75th percentiles; whiskers, 1.5 × IQR. (F) Heatmap is presented which illustrates the enrichment of transcription factor (TF) motifs at promoter (within 1 kilobase (kb) of the transcription start site (TSS)) and distal (more than 2.5 kb from the TSS) p300 peaks in L1C-specific, L2C-specific and shared regions. (G) The enrichment of Pol II, p300 and H3K27ac in p300 shared peaks-associated genes in the control (DMSO) and A485-treated embryos at the E2C and L2C stage is presented. TSS: transcription start site. TES: transcription end site. (H) Boxplots showing the normalized Pol II pausing index (PI) of p300 shared-peaks associated genes in control and A485 treatment embryos. Pol II PI was determined by odds ratio of read number mapped to promoter regions (-150 to +50 bp from TSS) and gene body regions (+50 bp downstream of TSS to transcription end site (TES)). P-value are indicated with two-sided Wilcoxon rank-sum test. Centre line, median; box, 25th and 75th percentiles; whiskers, 1.5 × IQR.

We then examined how the inhibition of p300/CBP during the minor ZGA stage leads to failed ZGA by analyzing the occupancy of p300 and Pol II in the A485 treatment group. The accumulation of p300 at L2C-specific regions, which should have been established at E2C, was compromised following A485 treatment (Figure 3D). These defects persisted until the L2C stage, although some accumulation of p300 was still observed from E2C to L2C after restoring catalytic activity (Figure 3D). The failure of p300 to enrich at L2C-specific targets resulted in decreased Pol II accumulation (Figure 3D), which in turn led to the downregulation of associated genes at the L2C stage (Figure 3E and S4D; including 707 significantly down-regulated vs. 106 significantly up-regulated genes). Furthermore, the inability of p300 to be recruited to L2C-specific sites was accompanied by abnormal retention of p300 and Pol II at L1C-specific targets (Figure 3D), as well as ectopic activation of relevant genes in L2C embryos treated with A485 (Figure 3E and S4D; including 1496 significantly up-regulated vs. 590 significantly down-regulated genes). These ectopically activated genes were enriched in pathways related to multicellular organism development, cell differentiation, and cell proliferation (Figure S2D), consistent with their CG-rich promoters. These observations indicate that p300/CBP-catalyzed acetylation promotes the pre-configuration of p300 and Pol II in E2C embryos.

Notably, the decreased L2C-specific p300 peaks with CG-low promoters were more likely to contain motifs recognized by ZGA-specific TFs such as DUX, OBOXs, RARs, NR5A2 and others (Figure 3F; Table S6), some of which are known to play critical roles in regulating ZGA. Conversely, the increased L1C-specific and unaltered stage-shared p300 peaks, featuring CG-high promoters, showed a greater tendency to contain motifs recognized by TFs like SPs and KLFs (Figure 3F; Table S6). These findings suggest that the transition of p300 and Pol II localization from the CG-high promoters of developmental genes to the CG-low promoters of ZGA genes during the ZGA stage is governed by TFs, and this process relies on the acetyltransferase function of p300/CBP.

Interestingly, genes associated with p300 stage-shared peaks exhibited downregulated expression at the E2C and L2C stages following A485 treatment (Figure 3E). The enrichment of p300 and Pol II in the promoter regions of these genes showed no significant difference between the DMSO and A485 groups (Figure 3G), likely due to the default loading of p300 and Pol II in these GC-rich promoter regions as early as the L1C stage (Figures 2F and 3D). These findings suggest an impact on Pol II elongation in the A485 group. To confirm the involvement of p300/CBP catalytic function in Pol II elongation activity for these genes, we compared the Pol II pausing index (PI; see Methods). As expected, the Pol II PI was significantly increased upon A485 treatment at the E2C and L2C stages, indicating impaired Pol II elongation (Figures 3H and S5A). Additionally, H3K27ac enrichment was reduced in these genes, despite no significant change in p300 enrichment (Figure 3G). These results suggest that insufficient H3K27ac negatively impacts Pol II elongation in these genes within the A485 group^15^, even though these genes with CG-high promoter exhibited default loading of p300 and Pol II as early as the L1C stage (Figure S5A). Our findings underscore the role of p300/CBP acetyltransferase activities in promoting Pol II elongation in ZGA genes.

### Dynamic landscape of p300-based super-enhancers during ZGA

SEs are clusters of enhancers characterized by the binding of high levels of master TFs and mediators, driving the high-level expression of genes encoding key regulators of cell identity^19,20^. Strong occupancy signals of p300 are utilized to predict SEs^31,32^. In this study, we identified SEs at the ZGA stage by ranking enhancers defined by distal p300 and H3K27ac signals (>2.5 kb from TSS) using the ROSE algorithm^19^. Accordingly, we identified a total of 788 L1C-specific, 612 E2C-specific, and 1156 L2C-specific SEs based on previously described criteria^19^, and categorized the remaining enhancers as typical enhancers (TEs; 12,000 in L1C, 7,247 in E2C, and 11,539 in L2C) (Figure 4A; Table S7; see Methods). As expected, SEs exhibited broader and stronger p300 and H3K27ac signals compared to TEs (Figures 4B and S6A). Consequently, the active genes near SEs (SE-genes) were generally expressed at higher levels than those near TEs (TE-genes) at the E2C and L2C stages (Figure S6B).

**Figure 4.**
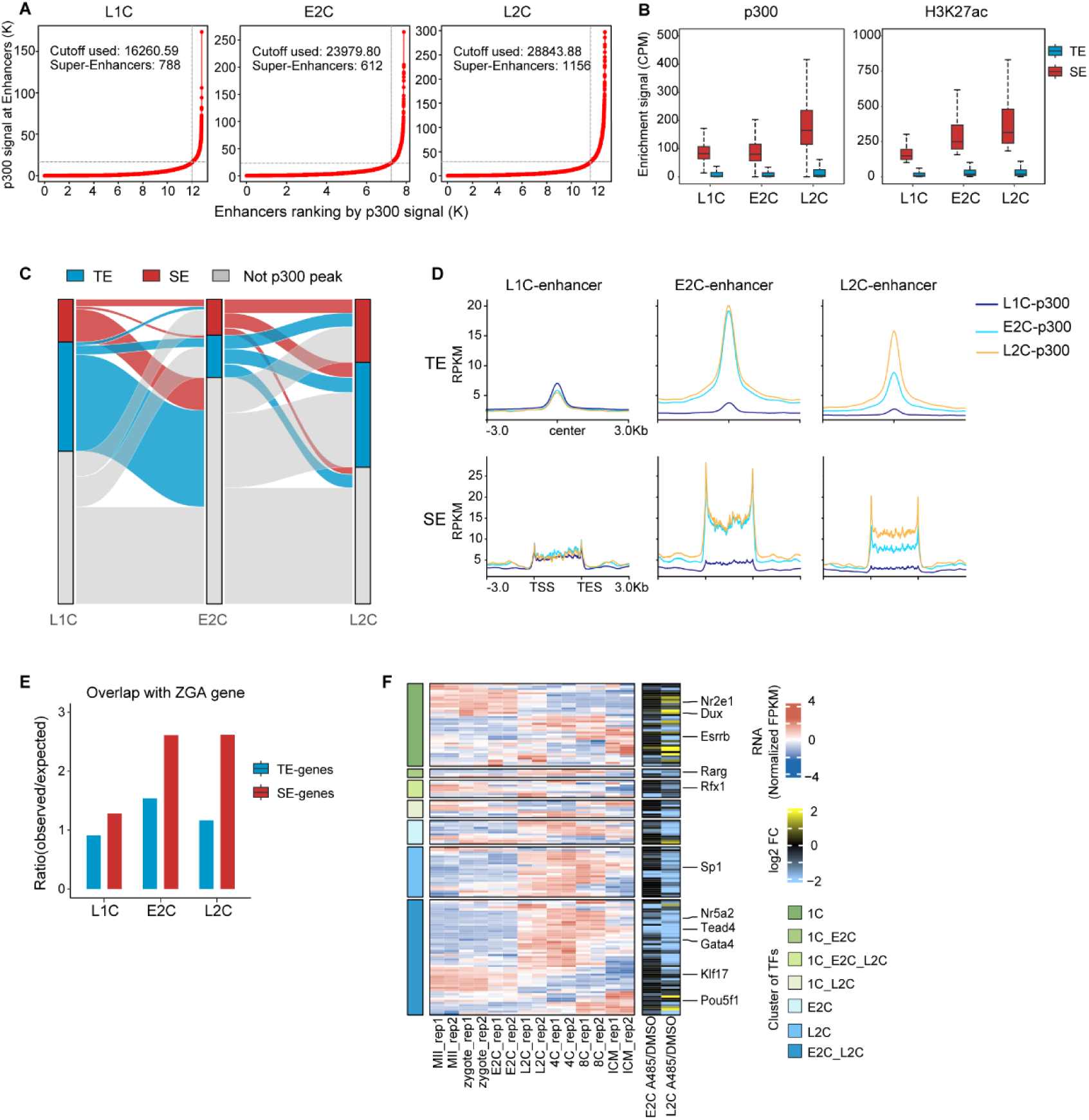
Dynamic landscape of p300-based super-enhancers. (A) Line graph illustrating the number of super enhancers (SEs) defined by the ranked p300 occupancy signal. (B) Boxplot illustrating a greater degree of p300 (left) and H3K27ac (right) enrichment in super enhancers (SEs) regions relative to typical enhancers (TEs) regions. Centre line, median; box, 25th and 75th percentiles; whiskers, 1.5 × IQR. (C) Sankey diagram illustrating the dynamic change in enhancer activities from L1C to L2C stages based on p300 signal. (D) Line chart showing the degree of p300 enrichment in typical enhancer (TEs) and super enhancer (SEs) regions from L1C to L2C stages. (E) Histogram illustrating the proportion of ZGA genes among SE- and TE-associated genes from L1C to L2C. (F) Heatmap showing the expression pattern of SE-associated transcription factors during mouse early embryonic development. The expression changes (A485/DMSO) at the E2C and L2C stages in response to A485 treatment are also shown.

We then observed dynamic changes in p300 enhancer landscapes from L1C to L2C. L1C-specific p300 enhancers nearly vanished in E2C embryos (Figures 3D and 4C). Meanwhile, L2C-specific p300 enhancers were established at E2C and persisted in p300 enrichment until L2C, differing from E2C-specific p300 enhancers, which were established at E2C and remained stable level throughout the 2C stage (Figures 4C and 4D). This observation suggests a progressive establishment of a distal p300-regulated enhancer activation program during ZGA stage. Notably, only 11.5% of enhancer-bound p300 is shared from L1C to L2C, in contrast to promoter-bound p300, where 33.4% of p300 peaks are shared across these three stages. These results indicate that enhancers during the ZGA stage are primarily activated starting at the E2C stage, unlike the inheritance of p300 at promoters from the L1C stage. DNA motif enrichment analysis revealed that OBOX motifs, known to play a critical role in ZGA, were the most frequent TF binding motifs in p300 distal peaks within both SEs and TEs at E2C and L2C embryos (Figure 3F; Table S6). In summary, we identified SEs and TEs in L1C, E2C, and L2C embryos, demonstrating the dynamic changes of p300 enhancer during the ZGA stage.

SEs are generally involved in key cell-type-specific gene expression, including master TFs and chromatin modification regulators^19^. Consequently, SE-genes exhibited stage-specific expression across the three developmental stages, conferring significantly higher transcriptional activity to ZGA genes in 2C embryos (Figure 4E). We then aimed to identify key TFs regulating ZGA genes by mapping SE-genes. The TFs associated with SEs were found to be stage-specific in L1C, E2C, and L2C embryos, aligning with their respective expression levels (Figure 4F). This set included genes known to play pivotal roles in ZGA and preimplantation embryo development, such as *Dux^8,10^* associated with L1C-SEs, *Nr5a2^12,33^* and *Yy1^34^*associated with L2C-SEs(Figure 4F; Table S6).

In addition to TFs, SE-genes were involved in epigenetic modifications (e.g., *Brd4*, *Kdm5b*, *Rbbp7* and *Ep300*)^26,35,36^ and various biological processes (e.g., *Nop2*)^37^ at the L2C stage (Table S6). Notably, p300 also regulated its own expression (Figure S2C; Table S2), suggesting a rapid positive feedback regulatory mechanism. Furthermore, several genes with uncharacterized functions in early embryo development were identified, warranting further investigation. Thus, our data reveal a complex regulatory network composed of SEs that govern genes implicated in ZGA and preimplantation embryo development.

### p300/CBP-catalyzed acetylation is critical for enhancer activities

To directly assess the role of p300/CBP activity in enhancer activation, we focused on the changes in distal p300 levels between the DMSO and A485 treatment groups. Notably, the frequency of distal p300 peaks significantly diminished in both L1C and E2C embryos following A485 treatment (Figure S3C). This indicates a tendency for p300 to accumulate in the promoter region in the absence of its acetyltransferase activity. This observation aligns with the erroneous persistence of high levels of promoter-bound p300 in L1C embryos treated with A485 (Figure 3D), which may be attributed to an enhanced chromatin binding affinity of p300 when its catalytic functions are inhibited^38^. In enhancer regions, p300 levels significantly decreased at the E2C and L2C stages following A485 treatment, accompanied by reductions in H3K27ac and chromatin accessibility (Figure 5A) at the E2C and L2C stages. Additionally, we analyzed the expression of enhancer RNAs (eRNAs), which are closely correlated with enhancer activities. The expression of eRNAs was dramatically reduced (Figure 5B), indicating that p300/CBP inhibition and decreased H3K27ac (Figure 5A) led to enhancer deactivation during the E2C and L2C stages. Concurrently, genes located near enhancers, particularly SE-genes that may harbor key determinants of cell identity, exhibited significant down-regulation (Figure S6C).

**Figure 5.**
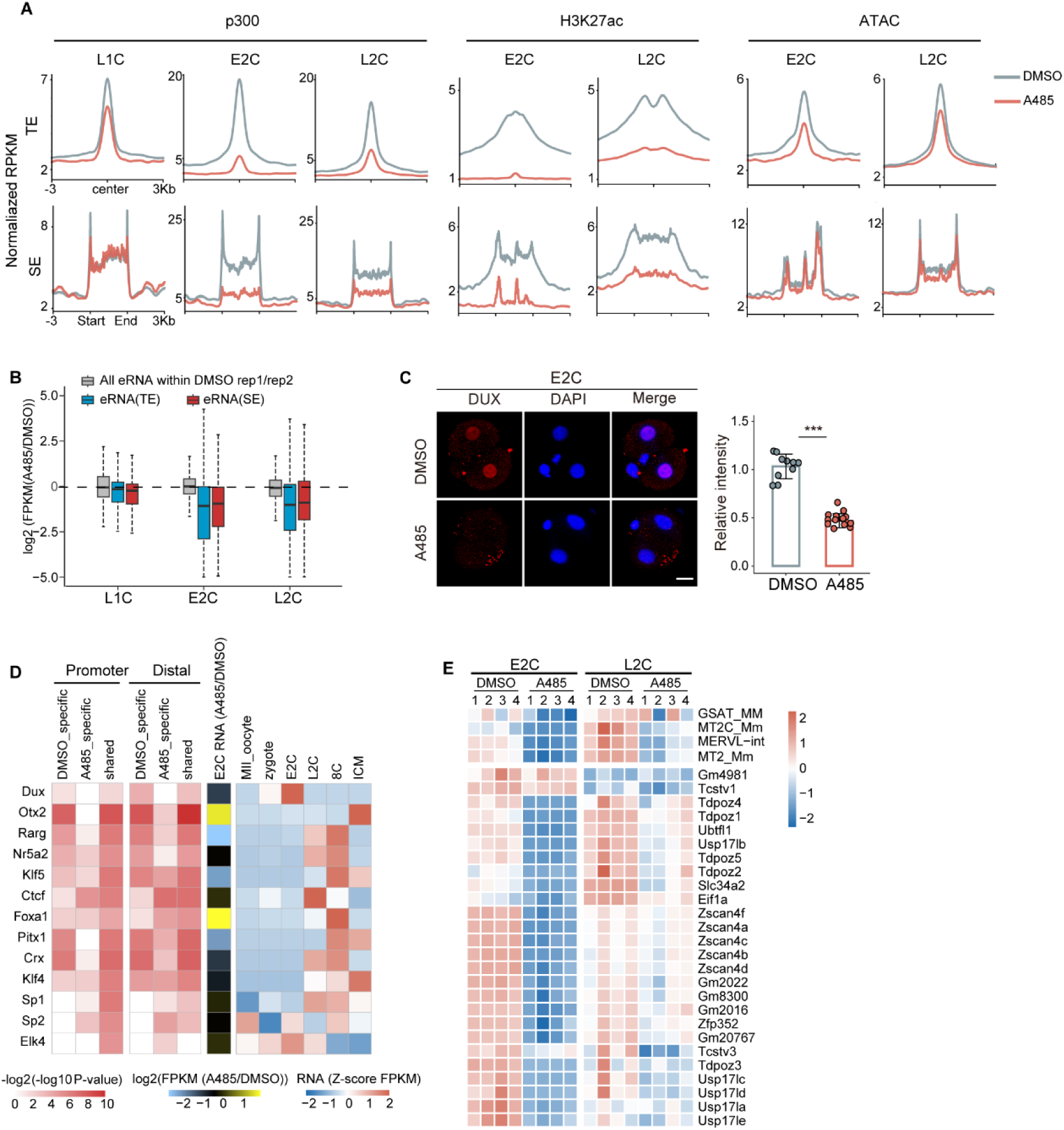
p300/CBP inhibition results in the inactivation of enhancer activities in mouse early embryos. (A) Line chart showing the degree of p300, H3K27ac enrichment and chromatin accessibility (ATAC) in typical enhancer (TEs) and super enhancer (SEs) regions in control (DMSO) and A485-treated embryos from L1C to L2C stages. (B) The expression changes of eRNA at typical enhancer (TEs) and super enhancer (SEs) regions after A485 treatment at L1C, E2C and L2C stages are presented. eRNA expression changes between DMSO biological replicates were calculated and shown as a control (gray). Centre line, median; box, 25th and 75th percentiles; whiskers, 1.5 × IQR. (C) Immunofluorescence images demonstrating the significant decrease of DUX signals in A485-treated embryos (n=18) compared with control embryos (DMSO-treated) (n=10) at the E2C stage. Scale bar: 20 μm. The quantification of DUX intensity is presented with mean S.E.M. *** P < 0.001 with two-sided t-test. (D) Left, the transcription factor (TF) motif enrichment at the promoter (within 1 kb of the transcription start site, or TSS) and distal (beyond 2.5 kb from the TSS) p300 peaks in regions that are specific to the DMSO treatment (DMSO-specific) (shared regions), those that are specific to A485 (A485-specific), and those that are shared between the two (Shared). Middle, the expression changes of transcription factors in response to A485 treatment at the E2C stage. Right, the expression pattern of transcription factors during the early stages of embryo development. (E) Heatmap illustrating the expression changes of DUX target genes in response to A485 treatment at the E2C and L2C stages.

Importantly, the expression of *Dux*, the TFs regulated by SEs in L1C (Figure 4F), was down-regulated in E2C embryos (Figures 1G and 5C), leading to a significant enrichment of the DUX motif in regions with significantly decreased p300 enrichment (Figure 5D; Table S8) and the decreased expression of DUX target genes (Figure 5E). These findings suggest that DUX acts as a master TF in E2C, governing enhancer formation and regulating the expression of key genes essential for the transition from minor to major ZGA. Our results indicate that p300/CBP activities are critical for enhancer functions in minor ZGA, which in turn regulate the key genes involved in initiating major ZGA.

### Exogenous *Dux* rescues the defects of ZGA and embryonic development induced by A485

In light of the impairment of minor ZGA associated with a deficiency of DUX (Figures 1G and 5C), which resulted in the failure of ZGA activation in the A485 treatment group, we sought to ascertain whether the exogenous *Dux*-overexpression (*Dux*-OE) might offer a potential avenue for the rescue of embryonic development. To this end, we injected *Dux-Flag* mRNAs into zygotes, followed by treatment with either DMSO or A485 (A485+*Dux* group; Figure 6A). Immunostaining assays confirmed the successful translation of the injected exogenous *Dux-Flag* mRNAs (Figure S7A).

**Figure 6.**
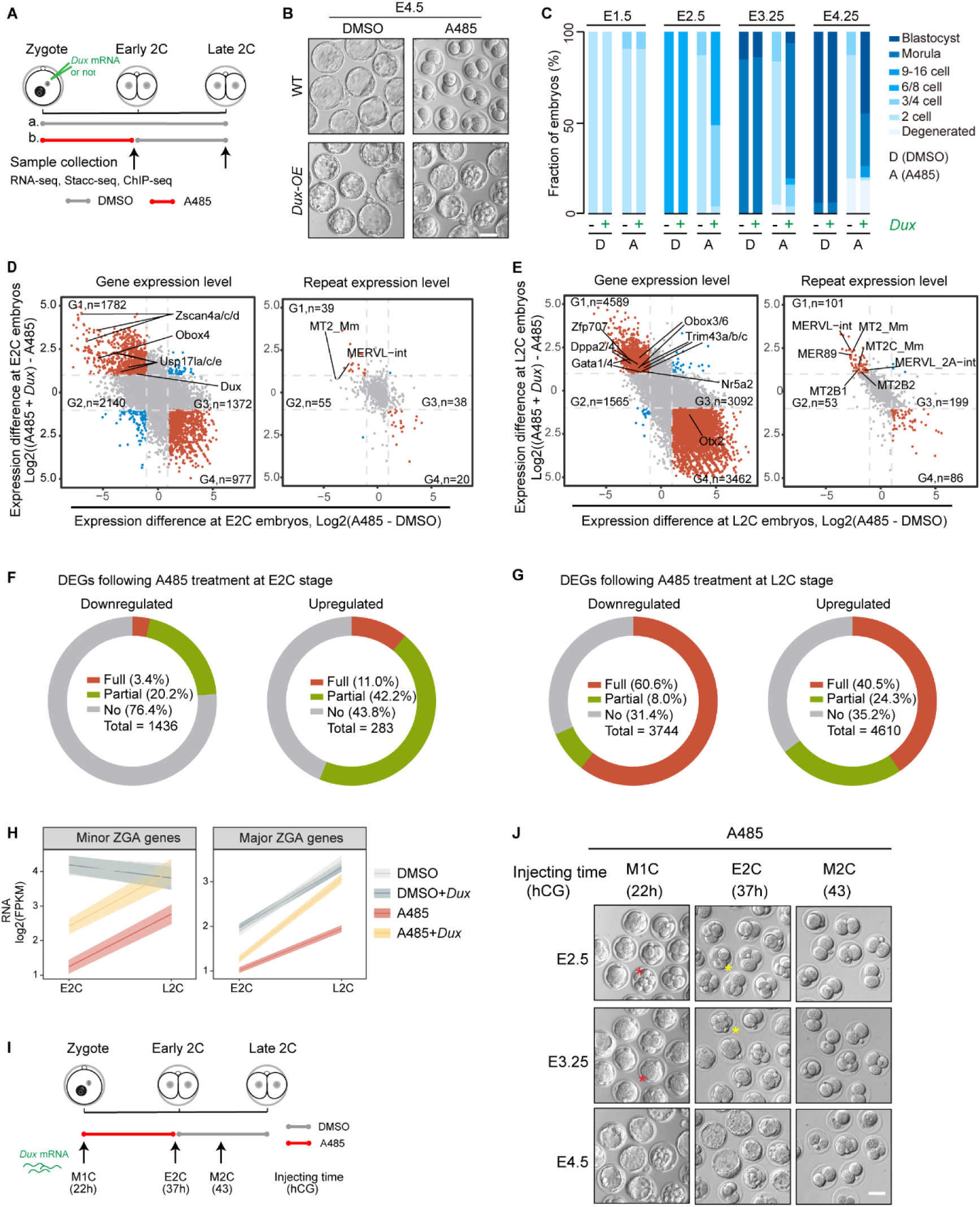
The defects caused by p300/CBP inhibition are rescued by overexpression of DUX. (A) Schematic illustrating the experimental procedure for the Dux-OE (Dux overexpression) experiment, which was conducted during the minor ZGA stage and involved the collection of RNA-seq, p300 Stacc-seq, Pol II Stacc-seq, H3K27ac ChIP-seq and ATAC-seq samples. (B) The image showing the embryo morphology of the control (DMSO), A485-treated (A485) and A485-treated with Dux overexpression (A485+Dux) embryos at E4.5. The injection of Dux-Flag mRNA was conducted at the middle 1-cell (M1C) stage. Scale bar: 50 μm. (C) The graph showing the developmental rate of the control (DMSO, D) and A485-treated (A485, A) embryos with (+) or without (-) *Dux* mRNA overexpression. (D) Scatterplots showing the rescue effect on genes (left) and repeats (right) expression in A485+Dux embryos at the E2C stage. The red dots represent genes and repeats whose expression is either upregulated or downregulated in A485 embryos in comparison to those treated with DMSO. In A485+Dux embryos, these genes and repeats are rescued. The blue dots represent genes whose expression is upregulated or downregulated to a greater extent in A485+Dux embryos. The dashed line represents the threshold of fold change (FC=2). (E) Scatterplots showing the rescue effect on genes (left) and repeats (right) in A485+Dux embryos at the L2C stage. The red dots represent genes and repeats whose expression is either upregulated or downregulated in A485 embryos in comparison to those treated with DMSO. In A485+Dux embryos, these genes and repeats are rescued. The blue dots represent genes whose expression is upregulated or downregulated to a greater extent in A485+Dux embryos. The dashed line represents the threshold of fold change (FC=2). (F) Donut charts illustrating the rescue of differentially expressed genes (DEGs) in A485+*Dux* embryos at the E2C stages, respectively. The term “full rescue genes” is used to describe those genes that are either downregulated or upregulated in A485 embryos (A485/DMSO) and subsequently upregulated or downregulated in the A485+*Dux* group to a significantly greater extent (A485+*Dux*/A485) (Padj < 0.05, FoldChange > 2 or < 0.5). The term “partial rescue genes” is used to describe those DEGs that are rescued in the A485+*Dux* group, but not to a statistically significant extent (Padj > 0.05, FoldChange >2 or <0.5; Padj < 0.05, FoldChange <2 or >0.5). The remaining downregulated and upregulated genes were not rescued. (G) Donut charts illustrating the rescue of differentially expressed genes (DEGs) in A485+Dux embryos at the L2C stages, respectively. (H) The expression changes of minor and major ZGA genes from the E2C to L2C stage in control (DMSO), A485, DMSO + *Dux* and A485+*Dux* embryos. The middle lines indicate the mean FPKM, while the background displays the 95% confidence interval around this value. (I) Schematic illustrating the experimental procedure for the Dux-OE (*Dux* overexpression) in M1C, E2C and M2C stages with A485-treatmet during minor ZGA stage. (J) The image showing the embryo morphology of A485-treated embryos with Dux overexpression in different time. Red asterisks represent normal development (8-cell at E2.5, morula at E3.5), while yellow asterisks represent delayed development by 0.5E (4-cell at E2.5, 8-cell at morula). Scale bar: 50 μm.

However, we observed that exogenous DUX-FLAG underwent significant degradation prior to the E2C stage (the A485 washing-off time point) after injection into zygotes (Figure S7A). This finding aligns with the critical role of DUX degradation in embryonic development^39^. Remarkably, in the A485 treatment group, nearly all embryos developed beyond the 2-cell stage, with approximately 50% and 30% progressing to the blastocyst and morula stages, respectively, following limited exogenous *Dux* mRNA injection (Figures 6B, 6C, and S7B). Furthermore, *Dux* exhibited a dosage-dependent effect, as excessive amounts led to a 2-cell arrest (Figure S7B). These results highlight the essential role of *Dux* in promoting embryonic development beyond the 2-cell stage following p300/CBP inhibition during the minor ZGA phase.

To understand the basis of developmental rescue following *Dux*-OE in the A485 group, we conducted transcriptome analysis (Figure 6A and S7C). At the L2C stage, embryos with *Dux*-OE demonstrated widespread rescue of DEGs in the A485 group, with 2,568 out of 3,744 (68.6%) of downregulated genes and 2,985 out of 4,610 (64.8%) of upregulated genes showing significant recovery (Figures 6E and 6G). This included a global restoration of major ZGA genes such as *Obox3/6*, *Nr5a2*, and *Yy1* (Figure 6E). Additionally, *Dux*-OE rescued the defective expression of *Kdm5b* and restored H3K4me3 modification at the L2C stage (Figure S7E).

At the E2C stage, *Dux*-OE partially ameliorated the gene expression abnormalities observed in the A485 group (Figure 6D), although only a small fraction of DEGs were fully rescued (49 out of 1346 (3.4%) of downregulated genes and 32 out of 283 (11.0%) of upregulated genes; Figure 6F). Notably, the majority of genes significantly upregulated by *Dux*-OE in the A485 group at E2C were enriched for minor ZGA genes (30 out of 52, 57.7%), including *Obox4*, *Zscan4s and Usp17s* (Figure 6D). Furthermore, 22 of these genes exhibited direct binding of the DUX motif at their promoters in mESCs^40^ (Figure S7D), suggesting that *Dux* activates its target minor ZGA genes at E2C independently of the acetyltransferase activities of p300/CBP. Interestingly, although only a small portion of both minor and major ZGA gene expression was restored at the E2C stage, their expression levels were reinstated to baseline levels observed in the DMSO group by the L2C stage (Figure 6H). This indicates that *Dux-*OE triggers the initiation of minor ZGA, enabling the sequential activation of both minor and major ZGA to be completed by the L2C stage, once p300/CBP function is restored form the E2C stage. This finding prompted us to investigate whether the timing of DUX-induced minor ZGA activation is critical for ZGA recovery by delaying *Dux*-OE until E2C or M2C (Figure 6I). Intriguingly, 48% of embryos with *Dux*-OE at E2C, coinciding with the A485 washing-off time, were able to develop to the blastocyst stage, albeit with a developmental delay of approximately half a day (0.5E) compared to embryos injected with *Dux* mRNA at the zygote (M1C) stage (Figures 6J and S7B). However, embryos with *Dux*-OE at the M2C stage remained arrested at the 2-cell stage (Figures 6I and S7B). In summary, our results suggest that the appropriate timing and dosage of *Dux*-OE can effectively rescue the minor ZGA deficits caused by p300/CBP inhibition, thereby facilitating major ZGA activation and promoting embryonic development.

### DUX recruits pol II to activate minor ZGA genes independently of p300

To elucidate the mechanism by which *Dux*-OE rescues ZGA in the A485 group, we examined the changes in p300 and Pol II enrichment (Figures 6A and S8A). At the L2C stage, the enrichment of Pol II in the A485+*Dux* group demonstrated a global recovery, with Pol II exhibiting decreased levels in the L1C-specific region and increased levels in the L2C-specific region, reaching levels comparable to those observed in the DMSO group (Figure 7A). This aligns with the correction of rescued ZGA genes and ectopically activated genes (Figures 6G and 6E). Interestingly, while there was some global recovery in p300 and H3K27ac levels, it did not correspond to the robust recovery of Pol II, particularly in major ZGA gene regions (Figures 7A and S8B). This suggests that the restoration of Pol II enrichment at ZGA genes is not solely dependent on p300. Notably, p300 and H3K27ac enrichment significantly increased in distal enhancer regions following *Dux*-OE in the A485 group (Figure S8C), accompanied by elevated eRNA expression (Figure S8D). These findings suggest that recovered p300 enhancer activities facilitate Pol II recruitment and elongation in ZGA genes at L2C, a process that is dependent on p300/CBP activity, consistent with previous studies^15^.

**Figure 7.**
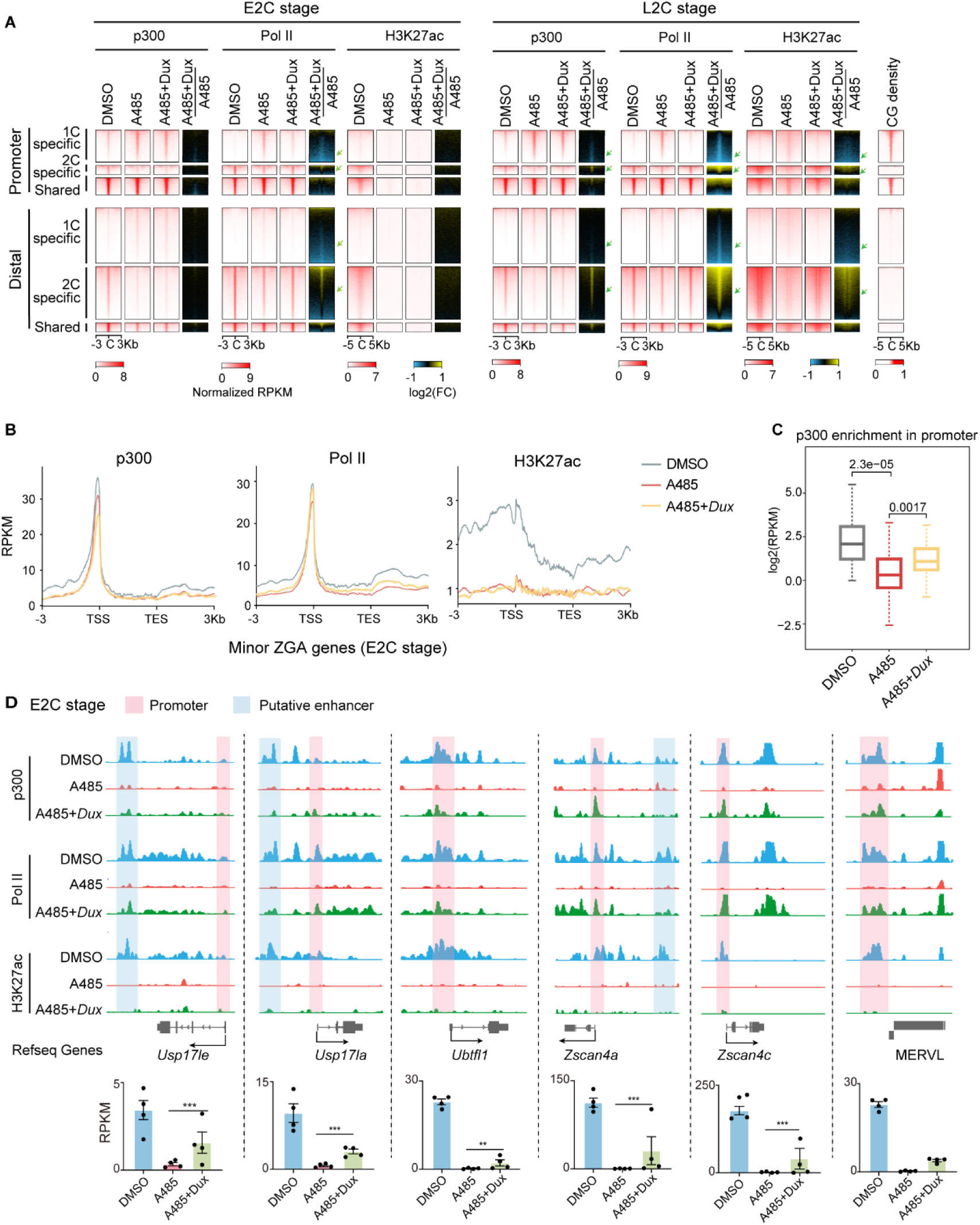
Dux recruits Pol II to activate minor ZGA genes with p300/CBP inhibition during the minor ZGA stage. (A) Heatmaps showing p300, Pol II and H3K27ac binding (Z-score normalized) at 1C-specific, E2C-specific, L2C-specific and shared p300 peaks in DMSO-control embryos (DMSO), A485-treated embryos (A485) and A485-treated with *Dux* injection embryos (A485+ *Dux*). Arrow, a significant recovery in A485+ *Dux* group. (B) Line chart showing p300, Pol II and H3K27ac enrichment at the minor ZGA gene region in E2C embryos. DMSO was used as the control. A485, A485-treated; A485+*Dux*, embryos treated with A485 and *Dux* overexpression. (C) Boxplot illustrating the p300 enrichment levels in promoter region (-150 to + 50 bp around TSS) in DMSO-control embryos (DMSO), A485-treated embryos (A485) and A485-treated with *Dux* injection embryos (A485+ *Dux*) at the E2C stage. The p-values are indicated using a two-sided Wilcoxon rank-sum test. Center line, median; box, 25th and 75th percentiles; whiskers, 1.5 × IQR. (D) The top panel presents IGV browser snapshots illustrating the enrichment of p300 and Pol II at DUX target genes in embryos treated with DMSO, A485, and A485+*Dux* at the E2C stage. The bar charts illustrate the gene expression patterns observed in the three aforementioned groups at the E2C stage. Data are mean ± S.E.M. *** *Padj* < 0.001.

At the E2C stage, there was a partial recovery of Pol II enrichment, particularly in minor ZGA genes (Figures 7A and 7B). This recovery was correlated with the increased expression in minor ZGA genes following *Dux*-OE (Figures 6D and 6F). Notably, the enrichment of p300 and H3K27ac did not exhibit a significant recovery, likely due to the inhibited catalytic activity of p300 at this time (Figures 7A and 7B). This observation further suggests that DUX can recruit Pol II and promote transcription to some extent independently of catalytic function of p300/CBP. Interestingly, despite the absence of restoration of p300 enrichment in all minor ZGA genes in A485+*Dux* embryos (Figure 7B), p300 levels were markedly elevated in the 22 *Dux* target minor genes that exhibited upregulation following *Dux*-OE (Figures 7C and S7D). This indicates that DUX can directly recruit p300^41^ to its target minor ZGA gene regions and activate their expression independently of p300/CBP’s catalytic function (Figure 7D). In conclusion, our findings reveal that *Dux* expression is contingent upon p300-mediated acetylation, whereas DUX is also capable of recruiting Pol II and activating downstream minor ZGA genes independently of p300/CBP’s acetylation function, facilitating the initiation of ZGA.

## Discussion

p300 and CBP are essential transcriptional co-activators that play a pivotal role in regulating transcription^15^. Our analysis of p300 occupancy in the genome during the ZGA stage reveals a dynamic pattern that is analogous to that observed for Pol II (Figures 2B and 8A). It is noteworthy that p300 plays a pivotal role in recruiting Pol II to the regions of ZGA genes. At the L1C stage, p300 initially binds to CG-rich promoter regions, associating indiscriminately with both developmental and ZGA genes. By the E2C stage, p300 predominantly enriches in the L2C-specific region, dissociating from developmental genes and binding to CG-poor promoters of major ZGA genes. This p300 enrichment is then maintained until the L2C stage. The acetylation target H3K27ac also exhibits pre-configuration prior to ZGA, with its distribution in E2C and L2C embryos displaying a comparable pattern^26^.

**Figure 8.**
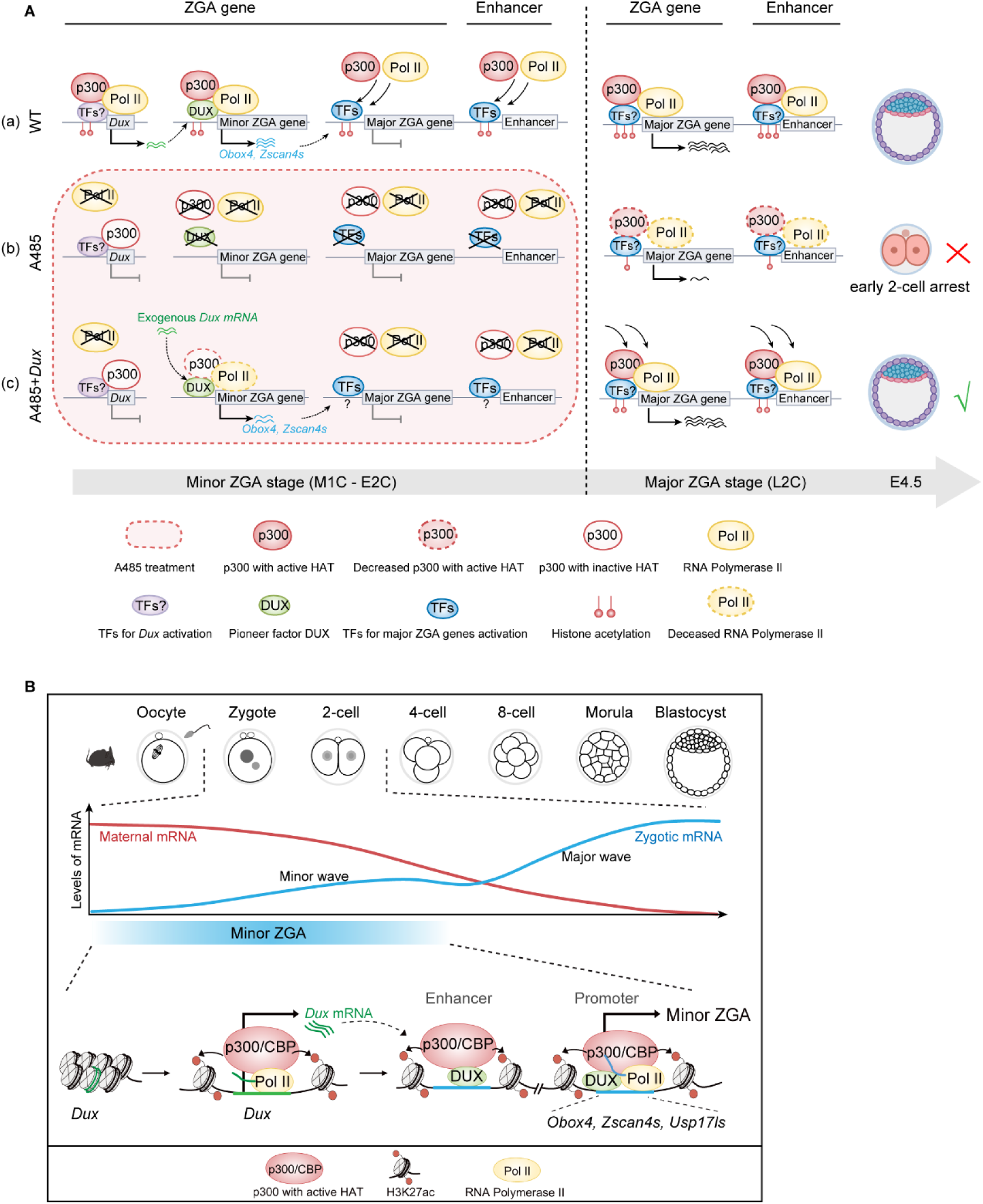
Model depicting the role of p300/CBP in mouse ZGA. (A) A model illustrating the role of p300/CBP in mouse ZGA. (a) Following fertilization, p300 initially binds to high-CG promoter regions, including developmental and some ZGA genes at the L1C stage. As transcription becomes active and progresses through one cell cycle to the E2C stage, p300 gradually dissociates from developmental gene regions. Under the influence of transcription factors, p300 then preconfigures in major ZGA genes with low-CG promoter and enhancer regions, thereby facilitating Pol II configuration and enhancer activation. During the major ZGA stage, p300 enrichment in enhancer regions persists, thereby enhancing enhancer activity and promoting the expression of major ZGA genes. (b) Inhibiting the acetyltransferase activity of p300/CBP during the minor ZGA stage leads to decreased chromatin accessibility, resulting in the failure of minor ZGA (down-regulated key genes including *Dux, Obox4, Zscan4*, etc.), which in turn impairs p300 and Pol II pre-configuration in the major ZGA gene and enhancer regions and causes abnormal retention of p300 in developmental gene regions at the E2C stage. This ultimately leads to major ZGA failure, even after restoring p300/CBP’s acetyltransferase activity at the E2C stage, resulting in embryonic development arrest at the 2-cell stage. (c) However, replenishing DUX during the minor ZGA stage enables DUX to recruit p300 and Pol II, thereby promoting the expression of minor ZGA genes such as *Obox4* and *Zscan4*. This subsequently leads to the enrichment of p300 and Pol II at major ZGA gene and enhancer regions during the L2C stage, although the precise mechanism remains unclear. As a result, the expression of major ZGA genes is restored, allowing embryos to develop to the blastocyst stage. (B) A simple model illustrating the role of p300/CBP in promoting minor ZGA via activating *Dux*. p300 and Pol II first accumulate near the *Dux* region, activating *Dux* transcription. Subsequently, *Dux* recognizes its target motif sequences, recruiting p300 and Pol II to activate the expression of minor ZGA genes.

The acetylation of H3K27ac, catalyzed by p300/CBP, is a hallmark of active enhancers^17^. p300 begins to establish itself at enhancer regions starting from the E2C stage, with its enrichment gradually increasing as development progresses to the L2C stage (Figures 3D and 4D). This suggests that the establishment of enhancer activity occurs progressively from the E2C to L2C stages, which is closely tied to Pol II elongation^15^. This may explain why ZGA is not immediately initiated after fertilization, despite the accumulation of transcription machinery at promoters in the 1C stage. Moreover, the gradual establishment of enhancer activities provides a critical time window for correcting Pol II transcription following fertilization ^3^. Meanwhile, SEs are distinguished by robust p300 signals, which facilitate the identification of the SE regulatory network and the prediction of potential master TFs and epigenetic regulators involved in ZGA and embryonic development (Figure 4F). Further investigation is required to ascertain the potential roles of these factors.

The acetyltransferase activity of p300 during minor ZGA is essential for the appropriate occurrence of ZGA. Inhibition of p300 results in the failure of p300 and Pol II configuration in ZGA gene regions, leading to stagnation at the initial developmental gene sites (Figure 3D). This disruption results in the downregulation of ZGA gene expression and the abnormal activation of developmental genes at the L2C stage (Figures 3E and S4D). Such dysregulation is consistent with the differential expression observed following the inhibition of HDAC1 deacetylase activity ^26,42^. It is noteworthy that the A485 treatment resulted in a reduction in HDAC1 expression at the L2C stage (Figure S2C). Furthermore, the absence of p300 acetyltransferase activity and reduced HDAC deacetylase activity results in a comparable pattern of differential H3K27ac enrichment^42^ (Figure S8E). An increase in H3K27ac enrichment in the p300 default binding region at the L1C stage indicates that p300 may facilitate the activation of HDAC, which erases H3K27ac to prevent the premature expression of developmental genes^42^.

The appropriate occurrence of minor ZGA is essential for the process of ZGA in mice and the early embryo development^3,43^. The loss of p300/CBP-catalyzed acetylation during minor ZGA results in the aberrant localization of p300 and Pol II at the E2C stage (Figure 3D). It is noteworthy that the absence of p300 acetyltransferase activity does not affect its own localization^15^, suggesting that p300/CBP-catalyzed acetylation during minor ZGA is crucial for regulating the expression of pivotal TFs or chromatin accessibility, which facilitate the pre-configuration of p300 and Pol II at the E2C stage. Our findings indicate that the inhibition of p300/CBP results in a notable reduction in the expression of the pivotal TF DUX. The overexpression of exogenous DUX at the zygote stage has been demonstrated to rescue the pre-configuration of Pol II at the E2C stage, thereby enabling embryos to develop to the blastocyst stage. It is noteworthy that exogenous DUX is degraded before the E2C stage, indicating that DUX downstream genes play a pivotal role in regulating Pol II localization and facilitating the transition from minor to major ZGA. Moreover, DUX activates these downstream genes independently of the catalytic function of p300/CBP, as p300/CBP remains inhibited at the E2C stage.

Furthermore, overexpression of DUX at the E2C stage, when p300 acetylation is restored, has been demonstrated to rescue development following A485 treatment, although this results in a delay of approximately half a day in embryonic development. This suggests that DUX is a crucial transcription factor for the initiation of ZGA. Prior research has demonstrated that *Dux* knockout (KO) in mouse embryos has minimal impact on the expression of a limited number of ZGA genes, with no discernible effect on embryonic development^39,44^. This may be attributed to the existence of additional genes that are functionally analogous to *Dux* and can nevertheless guarantee the occurrence of ZGA, such as *Obox4*^45^.

Furthermore, DUX plays a crucial role in facilitating the reprogramming of somatic cell nuclear transfer (SCNT) embryos^8,9^. The overexpression of DUX has been demonstrated to enhance the development of SCNT embryos to the blastocyst stage. In contrast, *Dux* homozygous knockout SCNT (*Dux*^-/-^) embryos have been observed to arrest at the 2-cell stage^9^. These findings provide further evidence that DUX functions as a pioneering transcription factor, promoting both ZGA and reprogramming. In SCNT embryos or those with impaired p300/CBP acetyltransferase activity, the expression of these crucial pioneer factors may be compromised. The overexpression of any one of these factors that are redundant with DUX could significantly enhance reprogramming. The roles of these key transcription factors warrant further investigation. In conclusion, our study presents compelling evidence that p300-catalyzed acetylation is a pivotal regulator of minor ZGA and elucidates the regulatory network of DUX in initiating ZGA in mouse embryos (Figure 8B).

## Limitation

This study primarily focused on investigating the role of p300/CBP acetylation activity in regulating ZGA using an inhibitor-based approach, leaving the structural functions of p300/CBP in ZGA regulation unclear. Additionally, we uncovered the crucial function of DUX in initiating minor ZGA by activating its target genes, even in the absence of p300/CBP acetyltransferase activity. However, the mechanisms underlying Dux activation remain unknown. Although we found that *Dux* expression is regulated by p300/CBP’s acetylation activity, it is possible that other maternal factors involved in *Dux* activation are yet to be discovered.

## Materials and Methods

### Embryo Collection and Culture

All experimental mice were raised and housed in the Laboratory Animal Center at Zhejiang University. All animal experiments were conducted in compliance with the guidelines for the care and use of laboratory animals and were approved by Zhejiang University. Female BDF1 mice (C57BL/6 × DBA2, Beijing Vital River Laboratory Animal Technology), aged 8-10 weeks, were super-ovulated with an injection of 7.5 IU of pregnant mare’s serum gonadotropin (PMSG) (San-Sheng Pharmaceutical, 110914564), followed by an injection of 7.5 IU of human chorionic gonadotropin (hCG) (San-Sheng Pharmaceutical, 110911282) 48 hours later. After hCG injection, females were mated with BDF1 stud male mice. Zygotes were collected from the oviduct 20 hours post-hCG, and the surrounding cumulus cells were removed using hyaluronidase (Sigma-Aldrich, H4272). Embryos were cultured in KSOM (Millipore, MR-016D) and incubated at 37°C with 5% CO2 in humidified air. Embryos were collected in M2 medium (Millipore, MR-015D) at the following time points after hCG: M1C (22 hours), L1C (30 hours), E2C (37 hours), M2C (43 hours), and L2C (48 hours).

### A485 treatment experiments

A485 (Selleck, S8740) dissolved in dimethyl sulfoxide (DMSO) (Sigma-Aldrich, D2650) was added to KSOM at a final concentration of 1 μM. To transiently inhibit p300/CBP, embryos were cultured in A485 at various time intervals post-hCG injection: 22 h∼37 h (minor ZGA), 37 h∼48 h (major ZGA), 22 h∼48 h (whole ZGA), 48∼116 h (from L2C to BL), 55 h∼116 h (from 4C to BL), 73 h∼116 h (from 8C to BL). Embryos were washed at least three times in KSOM to remove A485. In the control group, an equivalent amount of DMSO (0.1%) was added.

### In vitro transcription and microinjection

The plasmid pCW57.1-mDux-CA (Addgene, 99284) was used as a template to amplify the full-length mouse *Dux* for in vitro transcription. T7 promoter and Flag tag were added to *Dux* fragment. Mouse *Dux-Flag* mRNA was synthesized with the mMESSAGE mMACHINE T7 ULTRA Transcription Kit (Thermo Fisher, AM1345) following the manufacturer’s instructions. For DUX rescue experiments, *Dux-Flag* mRNA (15 ng/μl or 100 ng/μl) was injected into cytoplasm of M1C, E2C and M2C embryos (23 h, 37 h and 42h post hCG injection) with a Piezo-drill (Eppendorf) and Eppendorf Transferman 4r micromanipulators. Samples were injected at 5–10 pl per zygote or 2C embryo.

### Immunofluorescence

Samples were briefly washed with PBS, fixed with 4% paraformaldehyde (PFA) (Servicebio, G1101) for 15 min and permeabilized with 0.5% Triton X-100 in PBS for 30 min at room temperature (RT). Samples were then blocked in 10% fetal bovine serum (FBS) in PBS for 1 h at RT and incubated with primary antibodies for at least 1 h at RT or overnight at 4°C (p300, Abcam, ab275388; CBP, Abcam, 253202; FLAG, Sigma, F1804; H3K27ac, Active Motif, 39133; H3K18ac, Abcam, ab40888; H2BK5ac, Abcam, ab40886; H3K14ac, Abcam, ab52946; H4K16ac, Abcam, ab109463; H3K4me3, CST, C4208). The polyclonal DUX antibody was generated by HuaBio (Hangzhou, China) with peptides PQEEAGSTGMDTSSPSD^46^. After washing with 0.1% Triton X-100 in PBS, samples were incubated with secondary antibodies for at least 1 h at RT. Finally, the nuclei were stained with 4,6-diamidino-2-phenylindoley (DAPI) (Sigma-Aldrich, D9542) for 20 min at RT. Images were captured with a 40 × objective lens using Zeiss LSM880 confocal microscope system.

### RNA-seq library preparation and sequencing

For Smart-seq2, 50 blastomeres were used per reaction, and four replicates were performed for each group. The zona pellucidae of the embryos were removed with Acid Tyrode’s solution (Sigma, T1788). The Smart-seq2 procedure was performed as previously described with minor modifications^47^. Briefly, RNAs with a polyadenylated tail were captured, reverse transcribed and pre-amplified. After Tn5-mediated fragmentation, sequencing libraries were generated using the TruePrep DNA Library Prep Kit V2 for Illumina (Vazyme, TD502), following the manufacturer’s instructions. Paired-end 150-bp sequencing was performed on a NovaSeq (Illumina) at the Novogene Co., Ltd.

### Stacc–seq library generation and sequencing

For Stacc-seq, at least 200 blastomeres were used per reaction to obtain high-quality data for p300 (400 blastomeres) or Pol II (200 blastomeres), and two replicates were performed for each group. The Stacc–seq procedure was performed as previously described with minor modifications^3,5^. Briefly, 0.5 μl PA-Tn5 (Vazyme, S613) and 0.5 μg p300 antibody (Abcam, ab259330) or 0.5 μl PG-Tn5 (Vazyme, S612) and Pol II antibody (Active Motif 102660) were added to freshly prepared DB1 buffer (10 mM Tris-HCl pH 7.4, 150 mM NaCl, 0.5 mM spermidine, 2% glycerol, 1× EDTA-free with Roche complete protease inhibitor, 0.02% digitonin and 2 mM DTT) and incubated at 4 °C for 30 min (total volume 12.5 μl). Meanwhile, the zona pellucidae of the embryos were removed with Acid Tyrode’s solution (Sigma, T1788). Fresh samples were lysed in 37.5 μl DB1 buffer and rotated at 4 °C for 10 min. Subsequently, the sample was mixed with 12.5 μl of pre-incubated antibody, pG/A-Tn5 mixture, and 12.5 μl 5× TTBL (DB1 buffer with 50 mM MgCl_2_), followed by incubation at 37 °C for 30 min to cleave the chromatin around the binding sites. Finally, DNA purification and sequencing library preparation were carried out following the Stacc-seq protocol. Paired-end 150-bp sequencing was performed on a NovaSeq (Illumina) at the Novogene Co., Ltd.

### ULI-NChIP-seq library preparation and sequencing

For ULI-NChIP-seq, 300 blastomeres were used per reaction, and two replicates were performed for each group. The zona pellucidae of the embryos were removed with Acid Tyrode’s solution (Sigma, T1788). The ULI-NChIP-seq procedure was performed as previously described with minor modifications^48^. 500 ng of histone H3K27ac antibody (Active Motif, 39133) was used for each immunoprecipitation reaction. The sequencing libraries were generated using the NEB Next Ultra II DNA Library Prep Kit (NEB, 7645S), following the manufacturer’s instructions. Paired-end 150-bp sequencing was performed on a NovaSeq (Illumina) at the Novogene Co., Ltd.

### ATAC-seq library preparation and sequencing

For ATAC-seq, at least 200 blastomeres were used per reaction, and two replicates were performed for each group. The zona pellucidae of the embryos were removed with Acid Tyrode’s solution (Sigma, T1788). The ATAC-seq procedure was performed as previously described with minor modifications^49,50^. Fresh samples were lysed in lysis buffer with 0.05% digitonin at 4 °C for 10 min. Then, the samples were incubated with Tn5 transposase (Vazyme, TD502) to cleave accessible chromatin and add adaptors. After DNA purification, sequencing libraries were generated following the manufacturer’s instructions. Paired-end 150-bp sequencing was performed on a NovaSeq (Illumina) at the Novogene Co., Ltd.

## Data analyses

### RNA-seq data processing

Sequencing reads were aligned to the mouse genome (mm10) with Hisat2 (version 2.2.1)^51^ after removing adaptor sequences and low-quality reads by Trim_Galore(version 4.0; https://www.bioinformatics.babraham.ac.uk/projects/trim_galore/). The read counts and normalized FPKM values for each gene were calculated with featureCounts (version 2.0.1)^51^ and cufflinks (version 2.2.1)^52^. To quantify the repetitive sequences, TEtranscripts (version 2.2.3)^53^ was used to calculate the read counts based on RepeatMasker annotation from the UCSC genome annotation database. To calculate the read counts of *Obox4*, trimmed sequencing reads were also mapped to the reference *Obox4* mRNA sequence by Magic-BLAST^54^. The read counts were normalized by total reads and gene length for estimation of FPKM.

### Differential expression analysis and functional annotation

Differential expressed genes and repetitive sequences were identified by DESeq2 (version 1.42.1)^55^ with criteria set at adjusted *P* < 0.05, Fold Change > 2 or < 0.5. Gene Ontology terms of target genes were conducted with DAVID^56^ based on cellular components, molecular functions, and biological processes.

### Identification of minor ZGA genes, major ZGA genes, maternal genes and development genes

ZGA and maternal genes were identified with RNA-seq data from early mouse embryonic development^27^. Among them, genes with low expression in both GV and MII (FPKM < 5) and up-regulated expression in L1C or E2C (FPKM > 5, at least 3-fold) were minor ZGA genes, while genes with up-regulated expression in L2C (FPKM > 5, at least 3-fold up-regulated compared to GV/MII) were major ZGA genes. Genes that were highly expressed in GV/MII (FPKM > 5) but still up-regulated in L1C or L2C (FPKM > 5, at least 5-fold) were also recognized as ZGA genes. Maternal genes were defined as genes that were highly expressed in GV and MII (FPKM > 5), but down-regulated in L2C (at least 5-fold). The developmental genes were defined as PcG target genes in mESCs, of which the promoters (± 1 kb around the transcription starting sites (TSS)) are marked by RING1B and SUZ12^57^.

### Stacc–seq, ChIP-seq and ATAC-seq data processing

Stacc–seq, ChIP-seq and ATAC-seq sequencing reads were aligned to the mouse genome (mm10) with Bowtie2 (version 2.2.5)^58^ after removing adaptor sequences and low-quality reads by Trim_Galore (version 4.0). Low-mapping-quality (MapQ < 20) reads and PCR duplicates were filtered with Samtools (version 1.10)^59^ and Picard (version 2.23; https://broadinstitute.github.io/picard/). For downstream analysis, read counts were normalized to RPKM in a 100-bp window using ‘bamCoverage’ from deepTools (version 3.5.3)^60^ and the signal intensity was visualized with IGV. The RPKM was further normalized with Z-score transformation to minimize the batch effect. The correlation between replicates was calculated with Pearson correlation coefficient (RPKM for 2-kb bin across the genome). Heatmaps and profiles of signal enrichment were generated by computematrix from deepTools. To measure Pol II traveling ratio, the genes with gene body length > 1 kb were calculated for Pol II enrichment density in promoter region (-150 to + 50 bp around TSS) and gene body region (+ 50 bp downstream of TSS to transcription end site (TES)) with RPKM no less than 2, and odds ratios of the read densities of TSS region to read densities of gene body region.

### Peak calling and motif analysis

Stacc-Seq and ChIP-seq peaks were identified using MACS2 (version 2.2.7.1)^61^ with the following parameters: -- nolambda --nomodel. The peaks were annotated using ChIPseeker (version 1.32.1)^62^ based on RefSeq, UCSC and Ensemble databases. Promoters were defined as ± 1 kb around TSS, and peaks >2.5 kb away from TSS were defined as distal peaks. To compare the repetitive element enrichment, the number of overlaps between annotated repeats and Stacc-seq peaks or a randomly shuffled peaks was calculated as ‘observed/expected’ value. Stage specific peaks were obtained by BEDTools (version 2.2.7.1)^63^. Motif enrichment analysis of target peaks was performed using the findMotifsGenome.pl from Homer (version 4.11.1; http://homer.ucsd.edu/homer/motif/) with the following parameters: -size 200 -len 8, 10, 12.

### Identification of SEs and target genes

Enhancers were defined as the overlapped distal p300 peak and H3K27ac peak regions. Enhancers less than 12.5 kb away from each other were merged using the ROSE_main.py script in the ROSE program (http://younglab.wi.mit.edu/super_enhancer_code.html). The merged enhancers with p300 enrichment strength above the threshold were SEs, while the rest were TEs. The closest gene to the enhancer (the TSS closest to the SE) within ± 50 kb were defined as the enhancer-associated genes by ROSE_geneMapper.py script.

### eRNA expression analysis

To calculate read count of eRNA, the enhancer region defined by distal p300 and H3K27ac peaks were transformed into eRNA gtf files. With RNA-seq data, the read counts and normalized FPKM values for eRNA were calculated with cufflinks (version 2.2.1)^52^ based on the eRNA gtf files. For the accuracy of quantification, eRNAs in the gene body region and with low expression level (FPKM < 0.5) in wild-type group were excluded.

### Statistics

At least three biological replicates were performed for all experiments unless otherwise specified. Statistical analyses were conducted using GraphPad Prism 8 or R (https://www.r-project.org). Differences between two groups were assessed with two-tailed unpaired Student’s t-tests or two-sided Wilcoxon rank-sum. Immunofluorescence intensities were quantified using ImageJ software. *P* < 0.05 was considered statistically significant.

## Data availability

Previously published datasets used in the present work include RNA-seq of oocyte and early embryos (GSE71434)^27^, Pol II Stacc-seq of wild-type early embryos (GSE135457)^3^, H3K27ac CUT&RUN of wild-type early embryos (GSE207222)^26^, H3K27ac ChIP-seq of HDAC1 mutant L2C embryos (GSE182555)^42^, ATAC-seq of wild-type early embryos (GSE207222)^26^, p300 ChIP-seq of TSCs and ESCs (GSE110950)^32^, RING1B and SUZ12 ChIP-seq of ESCs (GSE119618)^57^. Our RNA-seq, Stacc-seq, ULI-NChIP-seq, and ATAC-seq data are deposited at GEO with accession no. GSE280521 and GSE280522.

## Acknowledgments

We are grateful to members of the Zhang laboratory for the discussion and comments during the preparation of the manuscript, and the Laboratory Animal Center at Zhejiang University for their support. This work was funded by the National Key R&D Program of China (2023YFD1300501), the National Natural Science Foundation of China (32161143032, 32072731 and 32472902).

## Author contributions

L.X. and K.Z. conceived the project and designed research; L.X., Y.D., P.Z. and S.L. performed the experiments; L.X. and H.J. analyzed the data; L.X., H.J. and K.Z. wrote the paper. Y.D., Y.S. and S.W. revised the manuscript.

## Competing interests

Authors declare that they have no competing interests.

## Notes

### Competing Interest Statement

The authors have declared no competing interest.

