## Supplemental information for "p300/CBP-catalyzed acetylation safeguards minor zygotic genome activation via activating DUX"

**Affiliations and Notes:** [1]Laboratory of Mammalian Molecular Embryology, College of Animal Sciences, Zhejiang University, Hangzhou, Zhejiang 310058, China; [2]L.X. and H.J. contributed equally to this work; [3]Present address: Center for Stem Cell Biology and Regenerative Medicine, MOE Key Laboratory of Bioinformatics, New Cornerstone Science Laboratory, School of Life Sciences, Tsinghua University, Beijing, China; [4]Present address: Assisted Reproduction Unit, Department of Obstetrics and Gynecology, Sir Run Shaw Hospital, Zhejiang University, School of Medicine, Hangzhou, Zhejiang, China; [5]Present address: Department of Histology and Embryology, School of Basic Medical College, Xinjiang Medical University, Urumqi, Xinjiang, China; [6]Present address: Obstetrics and Gynecology Hospital, Institute of Reproduction and Development, Fudan University, Shanghai, China; [7]Lead contact; [*]Correspondence:

**Keywords:** ZGA; p300/CBP; RNA Polymerase II; Enhancer; Preimplantation; Embryo

Supplementary Figure 1-8

Supplementary Table legend 1-8

Figure S1

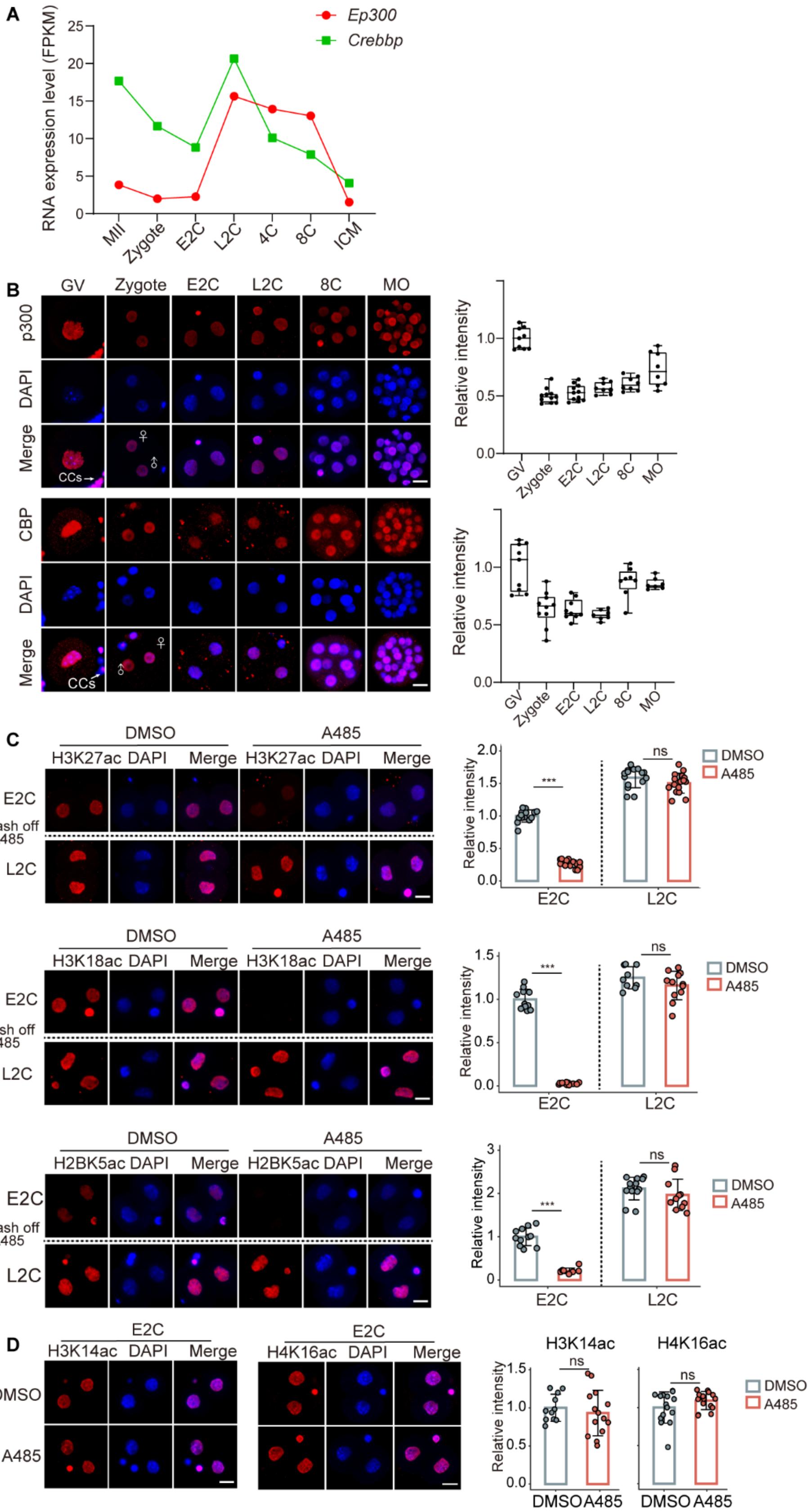


**Figure S1. A485 specifically inhibits the acetyltransferase activity of p300/CBP.**

(A) Line plots showing dynamic expression of *Ep300* and *Crebbp* in germinal vesicle (GV) oocytes and early embryos.

(B) Immunofluorescence showing p300 signal in mouse germinal vesicle (GV) oocytes (n=9), zygotes (n = 11), early 2-cell (E2C; n = 11), late 2C (L2C, n = 9), 8-cell (8C, n = 8) and morula (MO, n = 8) embryos and CBP signal in GV oocytes (n=9), zygotes (n = 10), early 2-cell (E2C; n = 9), late 2C (L2C, n = 7), 8-cell (8C, n = 8) and morula (MO, n = 7) embryos. Scale bar, 20 μm. CCs, Cumulus cells. Quantification of p300 and CBP intensity are shown in boxplots. Each dot represents a single embryo.

(C) Immunofluorescence showing histone acetylation signal catalyzed by p300/CBP in A485-treated embryos and control embryos (DMSO-treated) at E2C and L2C stage. H3K27ac, E2C (DMSO, n=14; A485, n=14), L2C (DMSO, n=15; A485, n=16); H3K18ac, E2C (DMSO, n=12; A485, n=11), L2C (DMSO, n=10; A485, n=13); H2BK5ac, E2C (DMSO, n=11; A485, n=9), L2C (DMSO, n=13; A485, n=11). Scale bar, 20 μm. Quantification of H3K27ac, H3K18ac and H2BK5ac intensity are shown with mean S.E.M. *** $P < 0.001$ with two-sided *t*-test. n.s., no significant difference with two-sided *t*-test.

(D) Immunofluorescence showing histone acetylation signal non-catalyzed by p300/CBP in A485-treated embryos and control embryos (DMSO-treated) at E2C stage. H3K14ac, DMSO, n=11; A485, n=15. H4K16ac, DMSO, n=15; A485, n=15. Scale bar, 20 μm. Quantification of H3K14ac and H4K16ac intensity are shown with mean S.E.M. n.s., no significant difference with two-sided *t*-test.

Figure S2

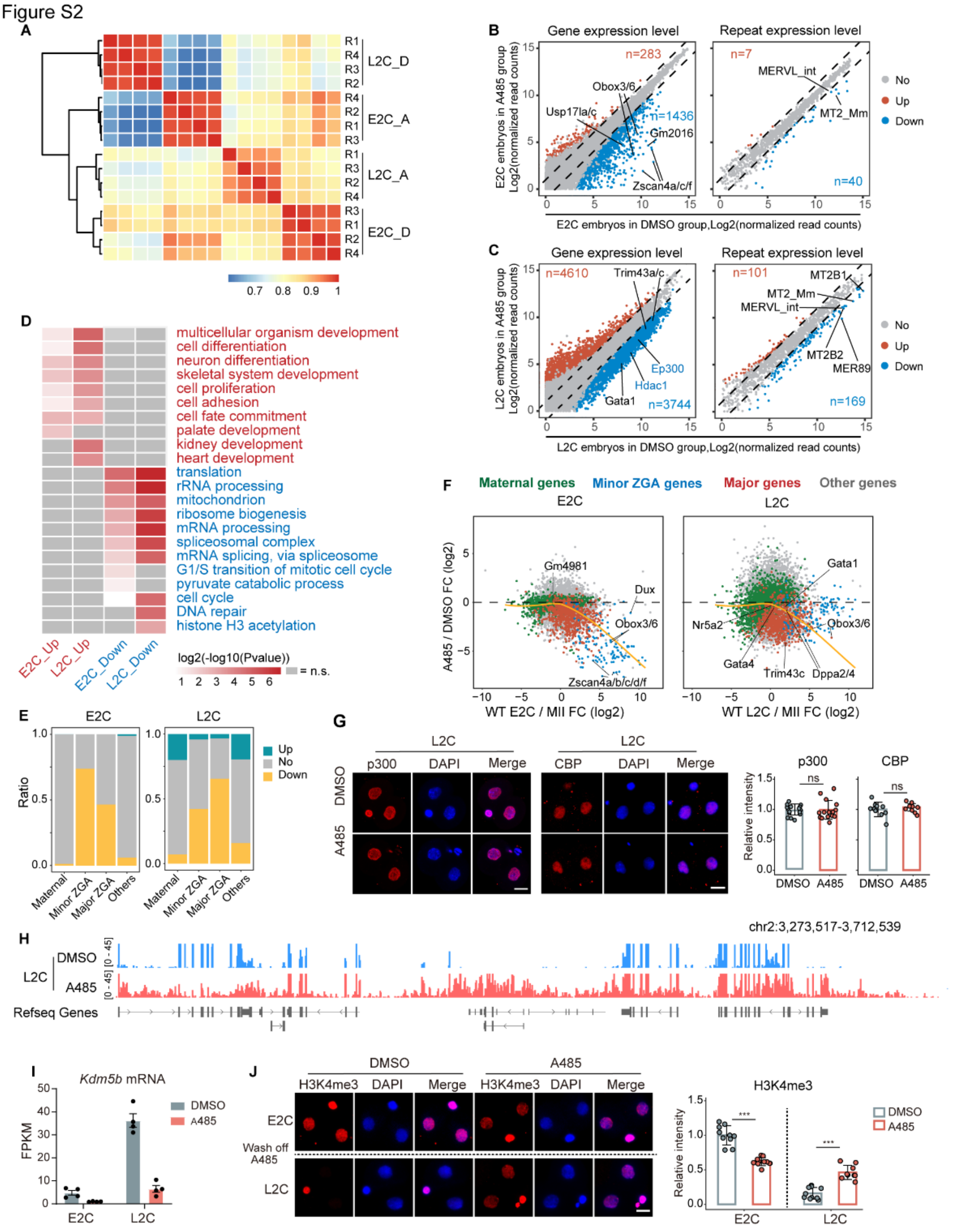


**Figure S2. p300/CBP inhibition during minor ZGA leads to minor and major ZGA failure.**

(A) Heatmap showing Pearson correlation between replicates among treatments and stages.

(B) Scatter plot showing the expression changes of genes (left) and repeats (right) upon A485 treatment in E2C embryos (4 biological replicates). Dashed line, the threshold of fold change (FC) (A485/DMSO). Differentially expressed genes (DEGs) are genes with FC > 2 or FC < 0.5, *Padj* < 0.05. Numbers of up- and down-regulated genes (red dots and blue dots, respectively) are indicated.

(C) Scatter plot showing the expression changes of genes (left) and repeats (right) upon A485 treatment in L2C embryos (4 biological replicates). Dashed line, the threshold of fold change (FC)

(A485/DMSO). Differentially expressed genes (DEGs) are genes with FC > 2 or FC < 0.5, *Padj* < 0.05. Numbers of up- and down-regulated genes (red dots and blue dots, respectively) are indicated.

(D) Heatmap showing the selected top Gene Ontology (GO) terms enriched among differential expressed genes (DEGs) identified in A485-treated embryos relative to control (DMSO) embryos at E2C and L2C stages. n.s., non-significant terms are shown in gray.

(E) Histograms showing the percentage of up-regulated (yellow), down-regulated (green), and unchanged (gray) genes in ZGA and maternal genes upon A485 treatment at the E2C and L2C stages.

(F) Scatter plot showing gene expression fold changes (FC) upon A485 treatment. Yellow lines, local regression fitting.

(G) Immunofluorescence showing p300 and CBP signal in A485-treated embryos and control embryos (DMSO-treated) at L2C stage. p300, DMSO, n=16; A485, n=15. CBP, DMSO, n=10; A485, n=9. Scale bar, 20 μm. Quantification of p300 and CBP intensity are shown with mean S.E.M. n.s., no significant difference with two-sided *t*-test.

(H) IGV browser snapshots showing gene expression in control (DMSO) and A485-treated embryos at L2C stage.

(I) Bar charts showing decreased *Kdm5b* mRNA in A485-treated embryos (4 biological replicates for E2C and L2C). Data are mean ± S.E.M. *** *Padj* < 0.001.

(J) Immunofluorescence showing H3K4me3 signal in A485-treated embryos and control embryos (DMSO-treated) at E2C and L2C stage. E2C (control, n=10; A485, n=11), L2C (DMSO, n=10; A485, n=9). Scale bar, 20 μm. Quantification of p300 and CBP intensity are shown with mean S.E.M. *** *P* < 0.001 with two-sided *t*-test.

Figure S3

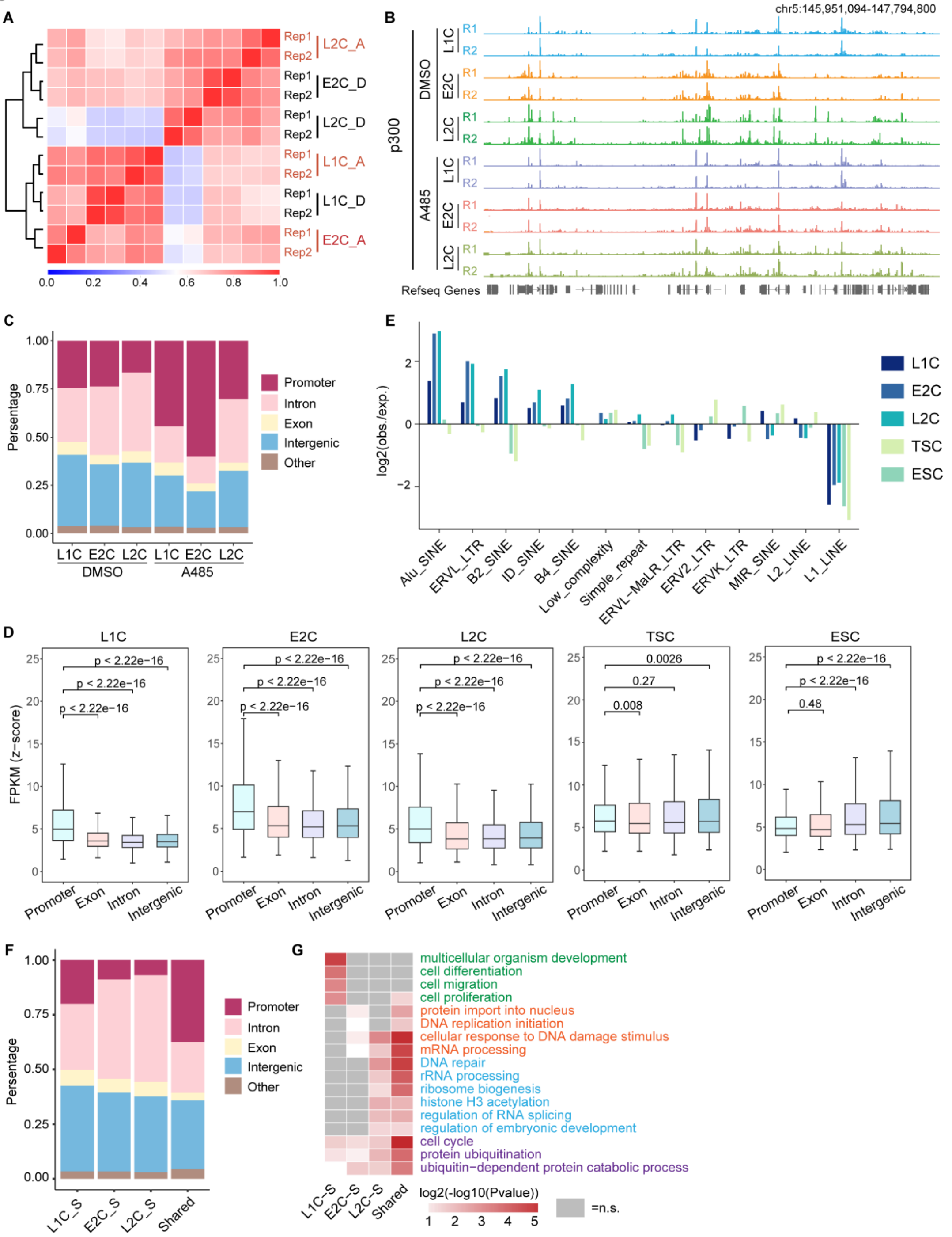


**Figure S3. The feature of p300-binding peaks during ZGA stage.**

(A) Heatmap showing Pearson correlation between the p300 signal in different treatment and stages.

(B) IGV browser snapshots showing p300 enrichment in control (DMSO) and A485-treated embryos at L1C, E2C and L2C stages (2 biological replicates for L1C, E2C and L2C).

(C) Bar chart showing the genomic distribution of p300-binding peaks in control (DMSO) and A485-treated embryos at L1C, E2C and L2C stages.

(D) Boxplot presenting the distribution of p300 occupancy scores across different genomic loci at L1C, E2C, L2C, TSCs (trophoblast stem cells)(Lee et al., 2019) and ESCs (embryonic stem cells)(Lee et al., 2019).

(E) Bar chart showing the repeat element enrichment at p300-binding peaks at L1C, E2C, L2C, TSCs (trophoblast stem cells)(Lee et al., 2019) and ESCs (embryonic stem cells)(Lee et al., 2019). obs., observed; exp., experimental.

(F) Bar chart showing the genomic distribution of p300-binding peaks in 1C-speific, E2C-specific, L2C-specific and shared p300-bindfing regions in control (DMSO) embryos.

(G) Heatmap showing the selected top Gene Ontology (GO) terms enriched among promoters p300 (around TSS < 1 kb) associated genes from 1C-speific, E2C-specific, L2C-specific and shared p300-binding regions. n.s., non-significant terms are shown in gray.

Figure S4

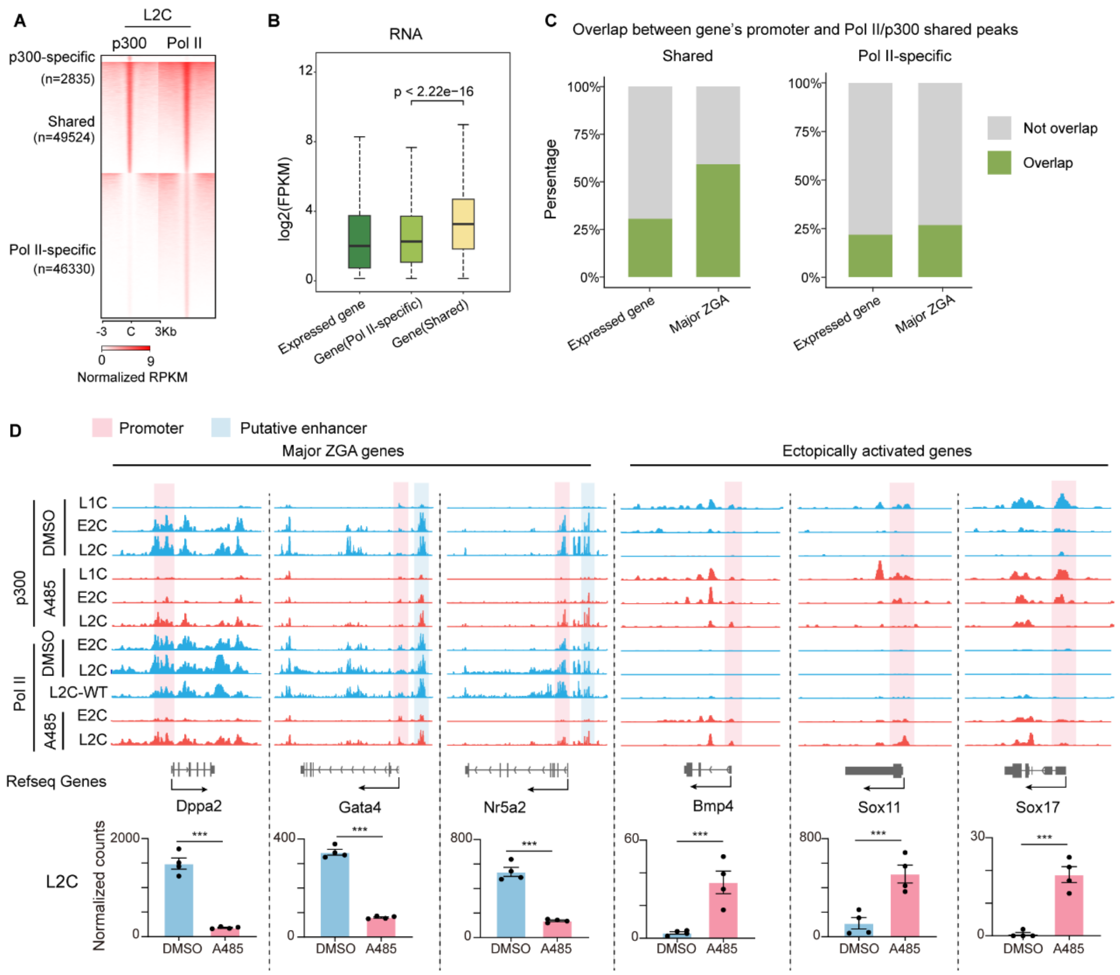


**Figure S4. A485 treatment resulted in pre-configuration defects of p300 and Pol II and failed Pol II elongation.**

(A) Heatmap showing the specific and shared enrichment of p300 and Pol II (GSE135457) at promoter regions in untreated L2C embryos.

(B) Boxplot showing the expression levels of genes associated with different types of promoter peaks in L2C embryos. P-value are indicated with two-sided Wilcoxon rank-sum test. Center line, median; box, 25th and 75th percentiles; whiskers, 1.5 × IQR.

(C) Histograms showing the proportion of expressed genes and major ZGA genes associated with shared-peaks (left) and Pol II-specific peaks (right) in L2C embryos. The proportion of expressed genes and major ZGA genes is accounted for by the overlapping and non-overlapping.

(D) Top, IGV browser snapshots showing p300 and Pol II enrichment at example genes in control (DMSO) and A485-treated embryos. L2C WT Pol II Stacc-seq dataset (GSE135457) were used. Bottom, bar charts showing gene expression between DMSO and A485 group. Data are mean ± S.E.M. *** *Padj* < 0.001.

Figure S5

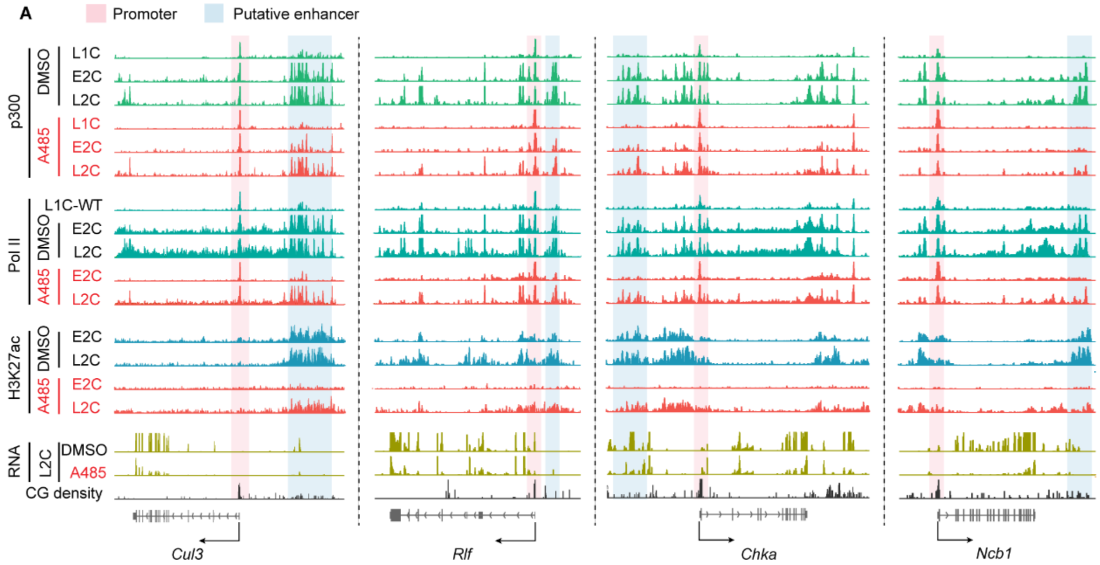


**Figure S5. The definition of super enhancers based on strong p300 signal.**

(A) IGV browser snapshots showing p300, H3K27ac and Pol II enrichment at genes associated with p300-shard peaks in control (DMSO) and A485-treated embryos. L1C WT Pol II Stacc-seq dataset (GSE135457) were used. RNA levels in DMSO and A485-treated embryos and CG density are shown.

Figure S6

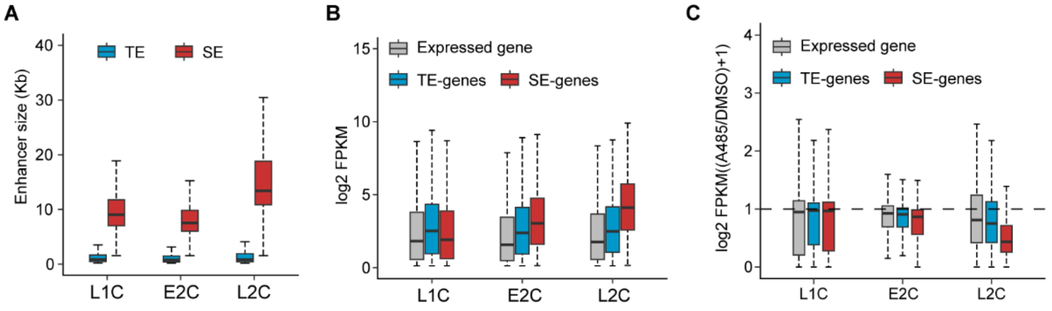


**Figure S6. The verification of super enhancers based on strong p300 signal.**

(A) Sequence lengths of typical enhancers (TEs) and super enhancers (SEs) at the L1C, E2C and L2C stages.

(B) Boxplot illustrating the expression levels of SE- and TE-associated genes at the L1C, E2C and L2C stages. Centre line, median; box, 25th and 75th percentiles; whiskers, 1.5 × IQR.

(C) Boxplot illustrating the expression change of SE- and TE-associated genes following A485 treatment at the L1C, E2C and L2C stages. Centre line, median; box, 25th and 75th percentiles; whiskers, 1.5 × IQR.

Figure S7

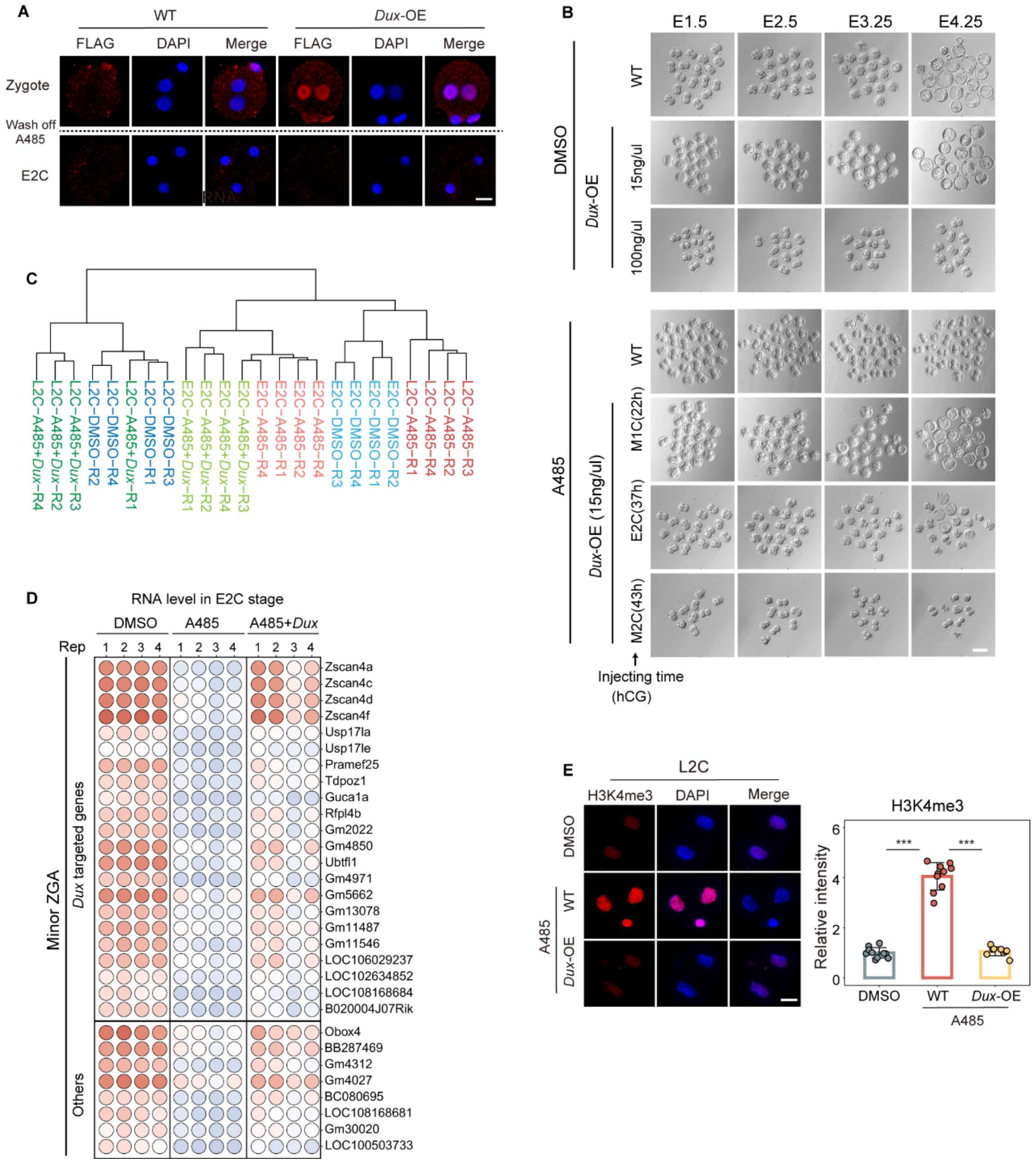


**Figure S7. Overexpressing *Dux* rescues the defects caused by p300/CBP inhibition.**

(A) Immunofluorescence showing FLAG signal in WT (wild type) embryos and *Dux*-OE embryos (*Dux* overexpression) at L1C and E2C stage. Scale bar, 20 µm.

(B) Picture showing the embryo morphology in DMSO, A485 and A485+*Dux* group during preimplantation development stage. Injecting *Dux-Flag* mRNA with different concentration and at different developmental stages individually (middle 1-cell (M1C), early 2-cell (E2C) and middle 2-cell (M2C)). Scale bar, 100 µm.

(C) Unsupervised hierarchical clustering of DMSO, A485 and A485+*Dux* embryos at E2C and L2C stages based on the FPKM in RNA-seq data (4 biological replicates).

(D) Balloon plot showing expression level of full rescued minor ZGA genes in A485+*Dux* embryos compared to A485 group. 22 genes whose promoter are bound by DUX directly are shown (4 biological replicates).

(E) Immunofluorescence showing H3K4me3 signal in A485-treated embryos (n=10), *Dux*-OE embryos (*Dux* overexpression with A485 treatment) (n=10) and control embryos (DMSO-treated) (n=9) at L2C stage. Scale bar, 20 μm. Quantification of H3K4me3 intensity are shown with mean S.E.M. *** $P < 0.001$ with two-sided *t*-test.

Figure S8

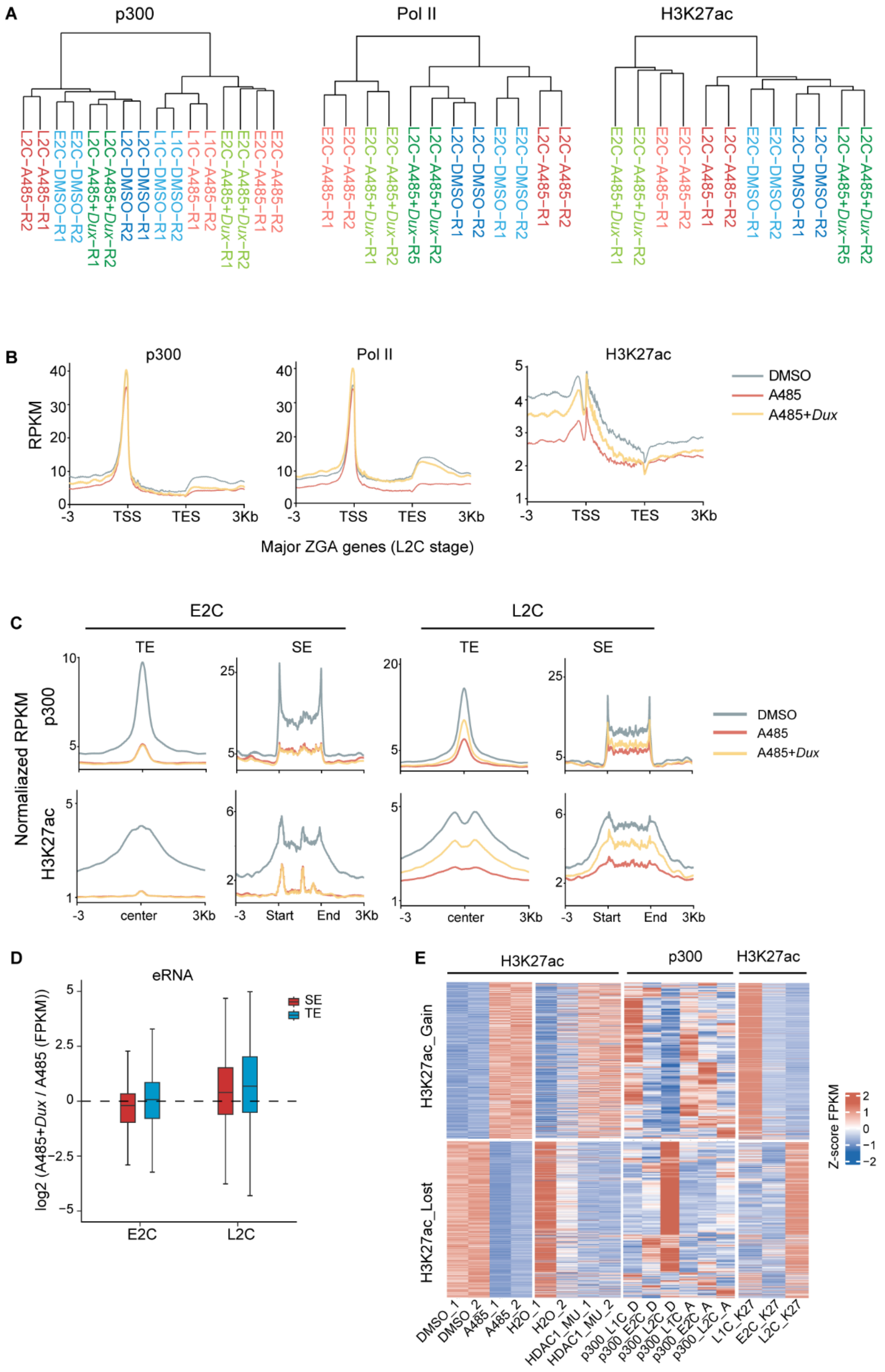


**Figure S8. Overexpressing DUX rescued the p300 and Pol II enrichment.**

(A) Unsupervised hierarchical clustering of p300, Pol II and H3K27ac signal (RPKM for 2-kb bin across the genome) in DMSO, A485 and A485 + *Dux* embryos at individual stages (2 biological replicates).

(B) Line chart showing p300, Pol II and H3K27ac enrichment at the major ZGA gene region in L2C embryos. DMSO was used as the control. A485, A485-treated. A485+*Dux* denotes embryos treated with A485 and *Dux* overexpression.

(C) The enrichment of p300 and H3K27ac in typical enhancers (TEs) and super enhancers (SEs) regions in DMSO, A485 and A485+*Dux* embryos at the E2C and L2C stage.

(D) The expression changes of enhancer RNAs (eRNAs) at typical enhancers (TEs) and super enhancers (SEs) in responsive to *Dux* overexpression in A485 group at E2C and L2C stages.

(E) Left, the H3K27ac enrichment changes between the DMSO and A485 group. H3K27ac enrichment shows the similar changes in the HDAC mutant group. Middle, p300 enrichment in the DMSO and A485 groups at the L1C, E2C and L2C stages. Right, H3K27ac enrichment in the wild-type L1C, E2C and L2C embryos. H3K27ac_Gain, regions where H3K27ac is significantly increased following A485 treatment at the L2C stage. H3K27ac_Lost, regions where H3K27ac is significantly decreased after A485 treatment at the L2C stage.

### Supplementary table legends

**Table S1. RNA levels (FPKM) in DMSO， A485 and A485+*Dux* embryos at the E2C and L2C stage.** Gene names and class for DMSO, A485 and A485+Dux embryos are included. 4 biological replicates.

**Table S2. Differentially expressed genes (DEGs) between DMSO and A485 embryos at the E2C and L2C stages.** Gene names, fold-changes and *Padj* at the E2C and L2C stages are included. *Padj* were calculated by DESeq2 with 4 biological replicates.

**Table S3. Differentially expressed genes (DEGs) between A485+*Dux* and A485 embryos at the E2C and L2C stages.** Gene names, fold-changes and *Padj* at the E2C and L2C stages are included. *Padj* were calculated by DESeq2 with 4 biological replicates.

**Table S4. p300 Stacc-seq peak in different classes in WT embryos.** Location, genomic region and group are included. The stage specific p300 peaks are identified by deeptools.

**Table S5. Stage specific p300 associated genes.** p300 associated genes in L1C-sepcific, E2C-specific, L2C-specific and shared region are included. p300 associated genes are identified by promoter p300 peak (TSS< 1 kb).

**Table S6. Motif enrichment of transcription factors in stage specific p300 peaks.** Motif enrichment and *P*-values for the transcription factors calculated by HOMER script findMotifsGenome.pl are included.

**Table S7. Super-enhancers and typical-enhancers identified by p300 peaks and their associated genes in L1C, E2C and L2C embryos.** Location, p300 enhancer associated genes and group are included. The genes with TSS overlapping the enhancer were defined as Overlap_GENES, the closest TSS as Closest_GENES, and those within ±50 kb of the enhancer as Proximal_GENES, all identified using the ROSE_geneMapper.py script.

**Table S8. Motif enrichment of transcription factors in decreased p300 peaks after A485 treatment at the E2C stage.** Motif enrichment and *P*-values for the transcription factors calculated by HOMER script findMotifsGenome.pl are included.
